# Extracellular cGAMP transmission exacerbates neuroinflammation in aging and multiple sclerosis

**DOI:** 10.64898/2026.09.30.754737

**Authors:** Dayanne R. Carvalho, Rina Wang, Nuo Liu, Yu Li, Talal Younis, Alessandra Sclip, Abinaya Muthusamy, Seula Shin, April Pawluk, Hani Goodarzi, Christoph Thaiss, Silvana Konermann, Lingyin Li

## Abstract

Neuroinflammation drives neurodegeneration and cognitive decline in aging, but the signals that spread this inflammation across the CNS remain unclear. Here, we show in mice that advanced age and experimental autoimmune encephalomyelitis (EAE) both result in elevated cytosolic mtDNA in microglia, which stimulates the production and export of cGAMP from these cells. Meanwhile, ENPP1, the dominant cGAMP hydrolase in the brain and spinal cord, is downregulated in both EAE mice and MS patients, releasing the brake on extracellular cGAMP accumulation. In Enpp1^H362A^ mice, which selectively lack cGAMP hydrolysis activity, unrestrained extracellular cGAMP worsens EAE-associated paralysis and drives premature aging phenotypes including impaired motor coordination (rotarod, pole test) and reduced exploration (open field test) in a manner dependent on STING. Single-cell RNA sequencing pinpoints microglia, astrocytes, and inhibitory neurons as the principal cGAMP responders that activate STING-dependent interferon-stimulated gene induction in response to elevated extracellular cGAMP. Mechanistically, microglia import cGAMP through the LRRC8A:C channel, while astrocytes and neurons import cGAMP via the glutamate antiporter system xc-, reinforced by SLC1A2-mediated glutamate uptake in astrocytes and SLC38A1/2-fueled glutamine-to-glutamate synthesis in neurons. In isolated primary astrocytes and neurons, high intracellular glutamate and system xc--driven redox demands favor cGAMP uptake, raising the possibility that neurotransmitter cycling and redox burden shape cellular vulnerability to cGAMP immunotransmission in vivo. Together, these findings position ENPP1 as a protective brake on neuroinflammation in aging and MS, nominating cGAS inhibition and CNS ENPP1 enhancement as therapeutic strategies.

## INTRODUCTION

The cGAS-STING pathway is an innate immune sensing mechanism in which the cytosolic DNA sensor cGAS produces the second messenger cyclic GMP-AMP (cGAMP), which then activates STING to drive type I interferon and inflammatory gene expression. While the cGAS-STING pathway is most often considered in the context of immunity and cancer through detection of cytosolic DNA originating from infection or genomic instability, there is mounting evidence that this pathway also plays a key role in neuroinflammation. Neuroinflammation is widely recognized as a central feature of neurodegeneration and age-related cognitive decline, and the cGAS-STING pathway plays pivotal roles in driving these diseases.^1–9^

In the neuroinflammatory context, microglia experience mitochondrial DNA leakage into the cytosol, which stimulates cGAMP synthesis and activates microglial STING signaling in a cell-autonomous manner, contributing to a proinflammatory microglial phenotype that exacerbates neuroinflammation.^7,9^ Neuronal STING signaling has been linked to senescence-associated phenotypes after ischemic stroke^10^ and has more recently been observed in neurons of multiple sclerosis (MS) patients and the EAE mouse model.^11^ However, neurons express low levels of cGAS, and neuronal cGAS has been shown to be dispensable for some neuroinflammatory-associated STING signaling,^11^ leaving the initiating signal for neuronal STING activation unclear.

cGAMP does not only act cell-autonomously: cancer cells and virally-infected cells are also known to export cGAMP into the extracellular space, where it serves as a paracrine signal.^12–15^ In cancer and infection contexts, extracellular cGAMP is internalized by infiltrating innate immune cells and stromal cells via cell type-specific transporters to activate STING signaling non-cell-autonomously.^14,16–22^ This intercellular cascade amplifies the effect of the original cGAMP signal and recruits and activates innate and adaptive immune cells at the site of cancer or infection. As a mechanism of immune evasion, many tumors express the cell-surface hydrolases ENPP1 and/or ENPP3 to deplete extracellular cGAMP and thereby attenuate the paracrine signal.^13–15,23,24^ It remains unknown whether and how extracellular cGAMP signaling occurs in the central nervous system (CNS), and how this paracrine cGAMP axis might contribute to STING-mediated neuroinflammation.

Here, we show that the CNS of mice experiencing aging or EAE exhibit elevated cGAMP levels, elevated STING signaling outputs, and a reduced ability to hydrolyze cGAMP compared to their young and healthy counterparts. MS patients showed similar reduction in ENPP1 expression levels. We demonstrate that ENPP1 is the dominant cGAMP hydrolase that restrains extracellular cGAMP signaling in the CNS, and that its expression is markedly reduced in aging and MS/EAE. Mice carrying an ENPP1 allele specifically deficient in cGAMP hydrolysis exhibit accelerated aging phenotypes including cognitive and motor deficits as early as 4 months of age, and experience exacerbated pathology following EAE induction. Microglia are major exporters of cGAMP, which enters other nearby microglia, astrocytes, and neurons through cell type-specific transporters to stimulate downstream STING signaling. Strikingly, we demonstrate that cGAMP can be imported by system xc- in exchange for the export of the neurotransmitter glutamate in neurons and astrocytes, uncovering a previously underappreciated connection between neurotransmission and neuroinflammation. Together, our work places the extracellular cGAMP axis, its checkpoint ENPP1, and the cell type-specific transporters that facilitate paracrine cGAMP communication as key mediators of neuroinflammation and aging in the CNS.

## RESULTS

### Extracellular cGAMP accumulation from ENPP1 deficiency exacerbates EAE

If extracellular cGAMP signaling operates in the CNS, we reasoned that the machinery to regulate it must also be present. To detect this activity, we first measured the rate of [^32^P]-cGAMP hydrolysis in tissue lysate from four brain regions and the spinal cord of wild-type (WT) C57/BL6 mice. While all brain regions displayed modest cGAMP hydrolysis activity, the spinal cord showed the most robust hydrolysis (Fig. 1A). Since the spinal cord is the primary site of pathology in EAE/MS, this finding raised the question of whether this cGAMP hydrolysis – and, by extension, cGAMP itself – plays a functional role in neuroinflammation.

**Figure 1.**
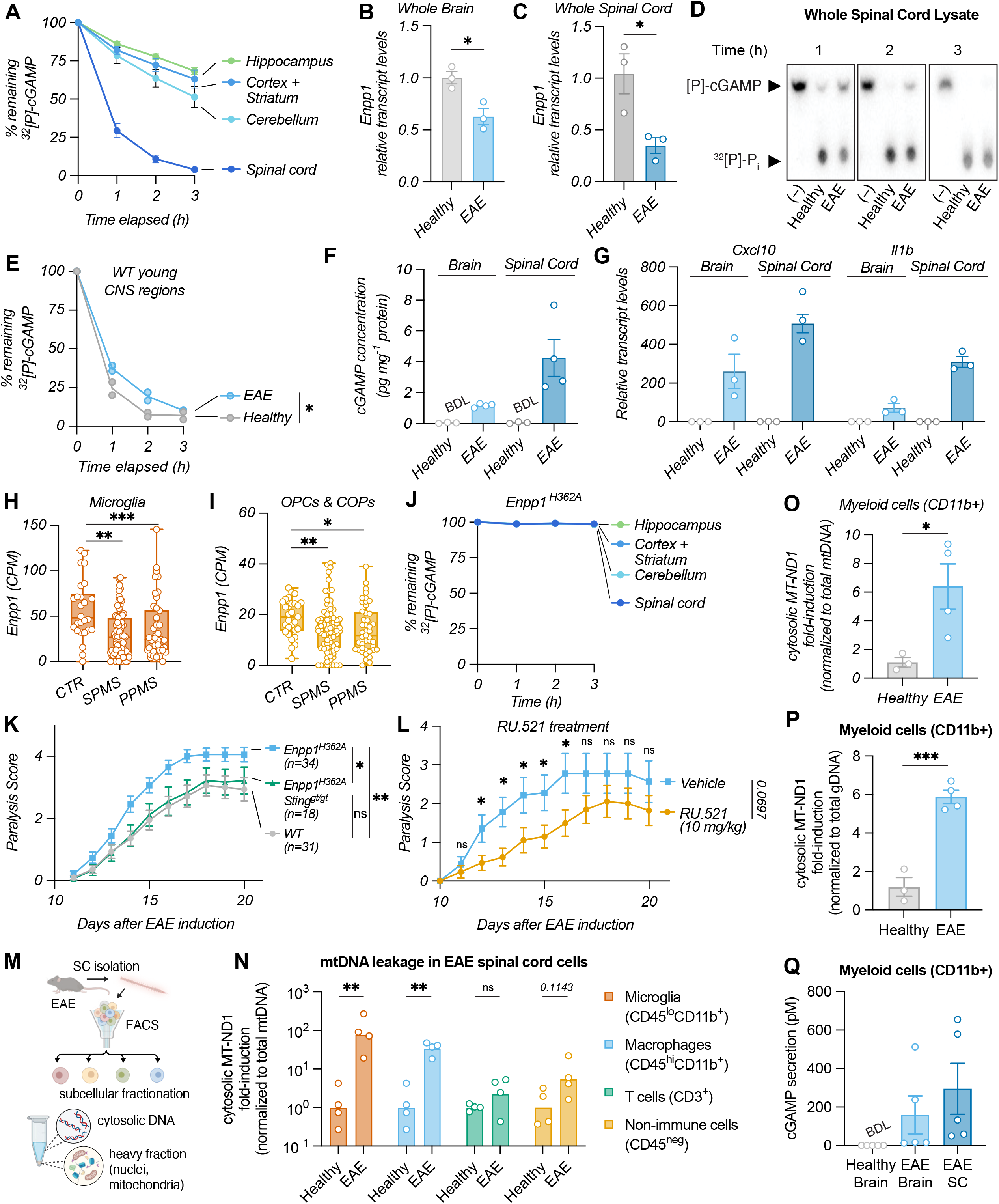
Extracellular cGAMP accumulation from ENPP1 deficiency exacerbates EAE. (A) Kinetics of 2′3′-^32^[P]-cGAMP degradation by tissue lysates from the indicated CNS regions of WT mice at 10 weeks of age (n=2 biological replicates). (B, C) mRNA expression levels of *Enpp1* in whole brain (B) and spinal cord (C) tissue of WT healthy (n=3 biological replicates) and MOG_35-55_-induced EAE mice at the symptomatic phase (paralysis score ≥ 2) (n=3 biological replicates). (D) Representative thin layer chromatography (TLC) tracking 2′3′-^32^[P]-cGAMP degradation by spinal cord lysate of WT healthy and EAE mice (paralysis score ≥ 2), at the indicated timepoints. (E) Kinetics of 2′3′-^32^[P]-cGAMP degradation by the spinal cord lysate of healthy and EAE mice (n=2 biological replicates). (F) 2′3′-cGAMP levels in healthy (n=3 biological replicates) and EAE (n=4 biological replicates) brain and spinal cord tissue measured by enzyme-linked immunosorbent assay (ELISA). (G) mRNA expression levels of *Cxcl10* and *Il1b* in whole brain and spinal cord tissue of WT healthy (n=3) and MOG_35-55_-induced EAE mice (n=3) mice. (H, I) Expression of *Enpp1* in microglia (H), oligodendrocyte precursor cells (OPCs), and committed oligodendrocyte precursors (COPs) (I) across the indicated MS diagnoses compared to healthy donors. Legends: **CTRL**: control; **SPMS**: secondary progressive MS; **PPMS**: primary progressive MS. (EGA: EGAD00001009169). Data are mean ± s.e.m. *p*-values were calculated using Mann-Whitney t-test. (J) Kinetics of 2′3′-^32^[P]-cGAMP degradation by tissue lysates of the indicated CNS regions of *Enpp1*^H362A^ mice at 10 weeks of age (n=2 biological replicates). (K) Paralysis scores for WT, *Enpp1*^H362A^, and *Enpp1*^H362A^ *Sting*^gt/gt^ mice at the indicated times after MOG_35-55_ immunization. Number of mice in each group is indicated in the legend. Significance calculated by two-way ANOVA. (I) Paralysis scores in WT mice treated with cGAS inhibitor RU.521 after EAE onset (n= 14 mice for vehicle and n=17 mice for RU.521 treatment). Data was pooled from 2 independent experiments. Significance of the treatment course was calculated by two-way ANOVA and significance of daily score was calculated by two-sided unpaired t-test with Welch’s correction. (M) Experimental scheme to measure mitochondrial DNA leakage in different cell types of the spinal cord during EAE symptomatic phase: spinal cord tissue single cell dissociation, followed by FACS, subcellular fractionation, and qPCR of mitochondrial genes. (N) Cytosolic mtDNA (mt-ND1) in CD11b^+^CD45^lo^ microglia, CD11b^+^CD45^hi^ macrophages, CD3^+^ T cells, CD45^−^ non-immune cells isolated from WT healthy and EAE mice, normalized to total mt-ND1 DNA in the heavy fraction (n=4 biological replicates per group). (O,P) Cytosolic mtDNA (mt-ND1) normalized to total MT-ND1 DNA (O) or total genomic DNA (B2M) (P) in the heavy fraction of MACS-sorted mouse primary spinal cord macrophages (CD11b^+^) from WT mice at 10 weeks of age, with or without MOG_35-55_-induced EAE (n=3-4 biological replicates per group). (Q) 2′3′-cGAMP secretion by MACS-sorted mouse primary brain and spinal cord macrophages (CD11b^+^) from wild-type mice with or without MOG_35-55_-induced EAE. Cells were cultured for 5 days after sorting and incubated for 48 h in the presence of ENPP1 inhibitor STF-1623. 2′3′-cGAMP levels in the supernatant were measured by enzyme-linked immunosorbent assay (ELISA) (n=5 biological replicates). Data are mean ± s.e.m. *p*-values were calculated using unless otherwise indicated.

Since ENPP1 has been reported as the dominant cGAMP hydrolase in the mouse CNS,^24^ we examined *Enpp1* expression in EAE mice. *Enpp1* transcripts were markedly reduced in both the brain and spinal cord of symptomatic 10-week-old mice ∼15 days post-EAE induction compared to healthy controls (Fig. 1B,C), which translated to decreased cGAMP degradation activity in whole spinal cord lysates as measured by [^32^P]-cGAMP hydrolysis (Fig. 1D,E). Interestingly, *Cgas* expression was also elevated in the spinal cord of EAE mice, while *Sting1* expression did not show a significant change between these groups (Fig. S1A,B). Consistent with increased cGAS expression and reduced ENPP1 activity, the spinal cord and brain of EAE mice displayed markedly elevated total cGAMP concentrations (Fig. 1F) and upregulation of the key type I interferon transcripts *Cxcl10* and *Il1b*, with the spinal cord showing a more pronounced effect in all cases (Fig. 1G).

To investigate whether these apparent shifts in extracellular cGAMP pathway components extend to human disease, we examined ENPP1 expression in post-mortem brain samples from patients with primary progressive (PPMS) or secondary progressive (SPMS) multiple sclerosis, from a published snRNA-seq dataset.^25^ The major CNS cell types expressing ENPP1 in the human brain are microglia, oligodendrocyte precursor cells (OPCs), and committed oligodendrocyte precursors (COPs) (Fig. 1H, I). Compared to healthy controls, we observed significantly lower ENPP1 transcript levels in the microglia, OPCs, and COPs from PPMS and SPMS patients (Fig. 1H, I). Beyond these cell types, neurons and oligodendrocytes express very low levels of ENPP1 in healthy controls (Fig. S1C-E); we observed no change in diseased neurons (Fig. S1C, D), whereas diseased oligodendrocytes exhibited an increase in ENPP1 expression (Fig. S1E). When we examined specific MS lesion types, *Enpp1* transcript levels were significantly lower in microglia from normal appearing white matter, active and chronic active demyelinated lesions, chronic inactive demyelinated lesions, and remyelinated lesions compared to healthy donors (Fig. S1F). Oligodendrocyte precursor cells from chronic active and -inactive demyelinated lesions also exhibited a decrease in *Enpp1* transcript levels (Fig. S1G), whereas excitatory neurons only showed this decrease in active demyelinated lesions (Fig. S1H). Meanwhile, inhibitory neurons and oligodendrocytes did not show significantly altered *Enpp1* levels in any type of lesion (Fig. S1I,J). While the mechanisms regulating ENPP1 expression in different cell types and lesion contexts remain unclear, these findings support the idea that a reduction in ENPP1-mediated extracellular cGAMP hydrolysis may be a common feature across mouse and human pathology in EAE and MS.

Given our observation of reduced ENPP1 levels and/or activity in both EAE and MS, we asked whether the capacity of the CNS to hydrolyze extracellular cGAMP has an impact on the pathology and progression of EAE. For these studies, we employed an *Enpp1^H362A^* mouse strain, in which ENPP1 is specifically deficient for cGAMP hydrolysis while retaining its other functions.^13^ The brain and spinal cord from these mice lack detectable cGAMP hydrolysis activity (Fig. 1J), confirming that ENPP1 is the primary contributor to this activity in the CNS.^24^ We induced EAE in three sets of mice: WT, *Enpp1*^H362A^ (lacking cGAMP hydrolysis), and *Enpp1*^H362A^ *Sting*^gt/gt^ (lacking both cGAMP hydrolysis and functional cGAMP-STING signaling). *Enpp1^H362A^* mice exhibited significantly exacerbated paralysis scores and higher disease incidence compared to WT controls, and these differences were abolished by the STING^gt/gt^ mutation (Fig. 1K, Fig. S1K). This initial result suggested that the loss of extracellular cGAMP hydrolysis – and the resultant accumulation of extracellular cGAMP – promotes neuroinflammatory phenotypes in a STING-dependent manner. Next, we asked whether the exacerbated pathology was dependent on continued cGAMP production by testing whether inhibiting cGAS *after* disease induction, at day 11, would improve the paralysis phenotype. Treatment with RU.521, a mouse cGAS-selective inhibitor, significantly slowed the progression of paralysis, even though treated animals still reached a similar peak disease severity as their untreated counterparts (Fig. 1L). This result suggests that continued cGAMP production accelerates disease progression, but its loss is not sufficient to prevent disease progression once it has begun.

Since mitochondrial dysfunction has been implicated in microglial-driven neuroinflammation in EAE and MS,^26^ and cytosolic mitochondrial DNA (mtDNA) is a known driver of cGAMP production, we asked whether myeloid cells contribute to the extracellular cGAMP pool. We isolated CD11b^+^CD45^lo^ microglia, CD11b^+^CD45^hi^ macrophages, CD3^+^ T cells, and CD45^−^ non-immune cells from the spinal cord of healthy and EAE mice. We used splenic T cells as healthy controls for the T cell population, given the negligible frequency of T cells in the healthy spinal cord (Fig. 1M). Following subcellular fractionation, we measured the amounts of cytosolic mtDNA by qPCR (Fig. 1N). Cytosolic mtDNA increased by ∼100-fold in microglia from EAE mice compared to healthy mice and by ∼35-fold in other myeloid cells, but did not significantly increase in encephalitogenic T cells (compared to healthy splenic T cells) nor in CNS resident non-immune cells. This signature of cytosolic mtDNA leakage persisted when CD11b⁺ myeloid cells isolated from EAE brains and spinal cord were cultured ex vivo for 1–2 weeks (Fig. 1O,P). Accordingly, these cultured myeloid cells from the EAE brain and spinal cord secreted high levels of extracellular cGAMP, which was undetectable in cultures from healthy CNS tissue (Fig. 1Q). These results demonstrate that, in the EAE context, elevated mtDNA leakage in microglia and other myeloid cells drives secretion of cGAMP into the extracellular space. Taken together with our earlier findings, this suggests a model in which this elevated extracellular cGAMP is insufficiently controlled by ENPP1 in EAE, exacerbating neuroinflammation and disease severity.

### ENPP1-mediated cGAMP hydrolysis is essential for healthy brain aging

Having observed that microglia export cGAMP in response to the acute disease state of EAE, we wondered whether microglia also export cGAMP in response to a chronic stress like aging, especially since aging microglia have elevated cytosolic mtDNA.^2^ Indeed, microglia isolated from aged mouse brains (18-25 months old) secreted cGAMP into the culture media at levels comparable to myeloid cells from EAE brains, while those isolated from young brains (2-3 months old) did not show detectable cGAMP export (Fig. 2A). To assess if this extracellular cGAMP axis also contributes to the chronic, low grade neuroinflammation found in the aging brain,^2^ we performed bulk RNA-sequencing to compare gene expression in aged (24-month-old) versus young (10-week-old) mouse forebrains (Fig. 2B). Consistent with prior reports of elevated STING signaling and increased expression of interferon-stimulated genes (ISGs) in the aging brain,^2^ we found that genes in the Interferon Alpha, Interferon Gamma, and Inflammatory categories were among the most upregulated pathways in aged forebrains. Gene Set Enrichment Analysis (GSEA) showed that the top three pathways significantly upregulated in the aged forebrains were downstream targets of cGAMP-STING signaling: Interferon Alpha Response, Interferon Gamma Response, and TNF-alpha Signaling via NF-κB (Fig. 2C). RT-qPCR confirmed upregulation of key ISGs in the brains (Fig. 2D) and spinal cords (Fig. 2E) of aged mice, with the brain exhibiting significant upregulation of all six genes tested whereas the spinal cord showed robust increases in 3 of 6 assayed transcripts.

**Figure 2.**
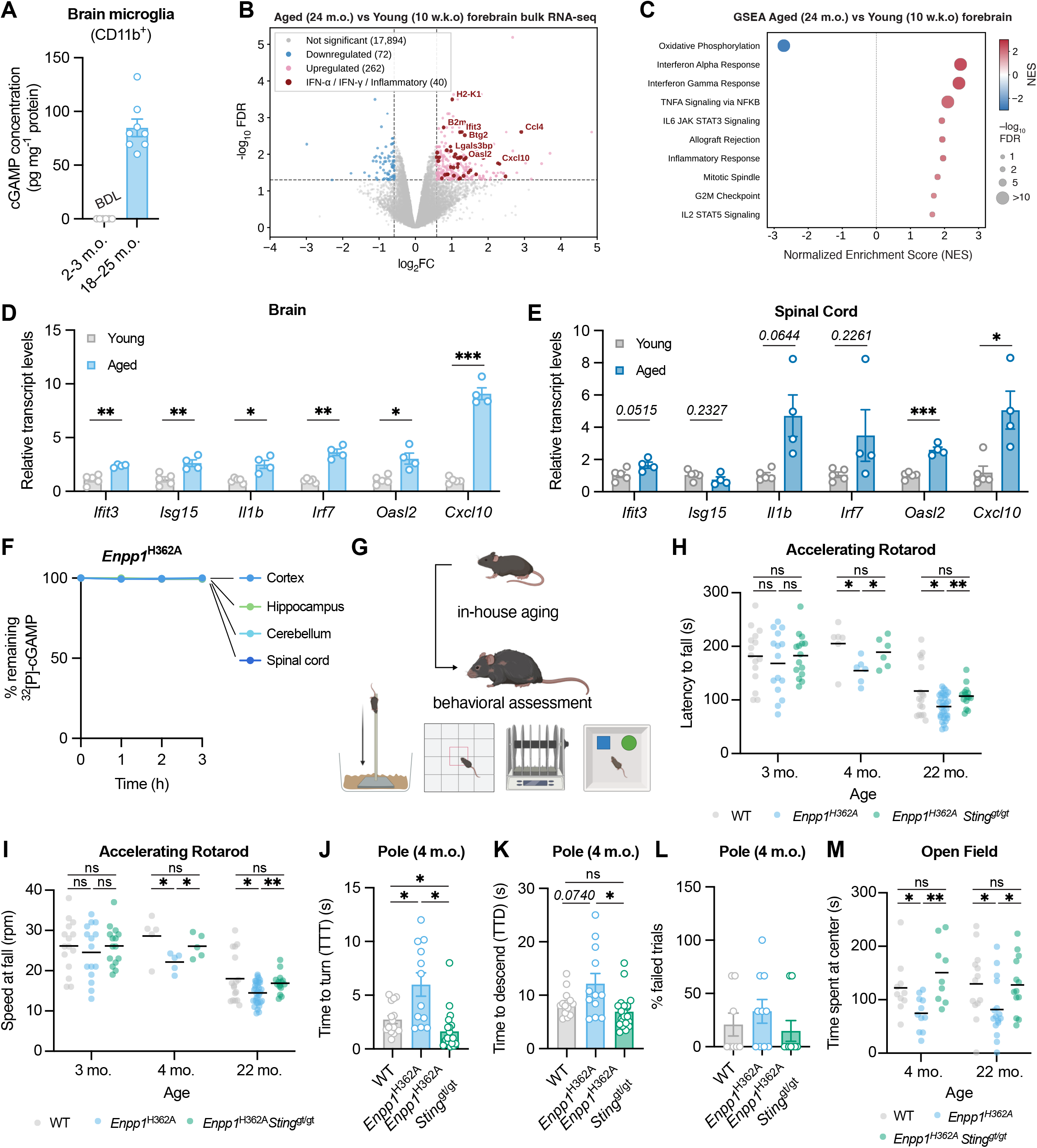
ENPP1-mediated cGAMP hydrolysis is essential for healthy brain aging. (A) MACS-sorted CD11b^+^ primary mouse brain microglia from aged wild-type mice were incubated for 48 h in the presence of ENPP1 inhibitor STF-1623. Supernatant was concentrated by solid-phase extraction with aminopropyl columns and 2′3′-cGAMP levels were measured by enzyme-linked immunosorbent assay (ELISA) and normalized to total protein amount in the cell lysates (n=8 biological replicates). (B) Differential gene expression from bulk RNA-seq analysis of young and aged mice forebrains (n=2 biological replicates per group). The volcano plot shows bolded type I IFN and inflammatory from the total list of DEGs (adjusted p-value ≤ 0.05, |[fold change (FC)]| ≥ 1.5). (C) Functional enrichment sorted on adjusted *p*-value in ascending order. Pathways identified with gene set enrichment analysis (GSEA) using the Hallmark gene set reveal interferon response as the top-enriched pathway in bulk RNA-seq analysis of young and aged mice forebrains (n=2 biological replicates per group). (D,E) Relative transcript levels of type I IFN and interferon-stimulated genes (ISGs) in whole brain tissue (D) and spinal cord tissue (E) of wild-type mice at 10 weeks of age (young) and 22–24 months of age (aged) (n=4–5 mice per group). (F) Kinetics of 2′3′-cGAMP degradation by different CNS regions of *Enpp1*^H362A^ mice at 22 months of age (n=2 biological replicates). (G) Strategy to assess the effect of extracellular cGAMP on aging-related neurodegeneration. (H, I) Average latency to fall (H) and average speed at fall (I) across three consecutive trials at the accelerating rotarod test with wild-type (n=6–17 per age group), *Enpp1*^H362A^ (n=6–32 per age group), and *Enpp1*^H362A^ *Sting*^gt/gt^ (n=6–15 per age group) (n=3–6 independent experiments performed for different age groups). (J,K,L) Time to turn (TTT), time to descend (TTD), and percentage of failed trials in the pole across three consecutive trials with wild-type (n=8), *Enpp1*^H362A^ (n=9 mice per age group), and *Enpp1*^H362A^ *Sting*^gt/gt^ (n=9 mice per age group) mice at 4–6 months of age (n=2 independent experiments). (M) Time spent at the center during open field testing of wild-type (n=10–13 per age group), *Enpp1*^H362A^ (n=11–16) and *Enpp1*^H362A^ *Sting*^gt^ (n=9–13) mice at 4 and 22 months of age. Data are mean ± s.e.m. *p*-values were calculated using two-sided unpaired t-test with Welch’s correction unless otherwise indicated.

Conversely, the oxidative phosphorylation (OXPHOS) pathway is the most significantly downregulated pathway, consistent with declining mitochondrial function with age. Mitochondrial dysfunction, with reduced OXPHOS as a key signature, triggers cytosolic mtDNA leakage, activates the cGAS-cGAMP-STING pathway and drives type I interferon and pro-inflammatory cytokine production, which further compromises mitochondrial function in a positive feedback loop.^27^ Together, these findings in the aged mouse CNS mirror our observations in the EAE model, raising the possibility that extracellular cGAMP may contribute to neuroinflammation in both acute and chronic contexts.

We next investigated whether extracellular cGAMP hydrolysis by ENPP1 regulates age-related neuroinflammation, as it did with EAE pathology. Brain regions from 22-month-old *Enpp1*^H362A^ mice displayed no detectable cGAMP hydrolysis activity (Fig. S2A, Fig. 2F), confirming that ENPP1 remains the dominant cGAMP hydrolase during aging with no detectable compensatory upregulation of alternative hydrolases. Consistent with an elevation in cGAMP concentration due to loss of its dominant hydrolase, RT-qPCR analysis of the whole forebrain (Fig. S2B) and spinal cord (Fig. S2C) from these aged mice showed elevated levels of ISG transcripts compared to their aged WT counterparts. Even at 4 months of age, *Enpp1*^H362A^ mice displayed elevated *Cxcl10* and *Il1β* expression in the hippocampus, striatum, and diencephalon compared to WT mice, but not in the other brain regions. Notably, these differences were abolished in *Enpp1*^H362A^*Sting*^gt/gt^ mice (Fig. S2D,E), confirming that the effect of extracellular cGAMP levels is dependent on downstream STING pathway activation. These findings suggest that extracellular cGAMP-STING signaling is active in several brain regions early in life and requires regulation by ENPP1 to protect against aberrant ISG induction.

We next examined the functional consequences of unregulated extracellular cGAMP–STING signaling across an aging time course by subjecting WT, *Enpp1*^H362A^, and *Enpp1*^H362A^ *Sting*^gt/gt^ mice to a series of motor and behavioral tests at different ages (Fig. 2G) using mice with similar weights (Fig. S2F). We first asked whether extracellular cGAMP affects motor coordination, grip, and strength by performing the rotarod test on 3-, 4-, and 22-month-old mice (Fig. 2H, I), using mice with similar Comparing WT mice across these ages, we observed that motor coordination improves from 3 months to 4 months of age, suggesting the acquisition of motor skills over time, as expected. WT mice performed much worse at 22 months compared to earlier time points, in agreement with previous studies that showed the decline of rotarod performance at advanced age.^28,29^ At 3 months, we did not observe any differences across the three genotypes, suggesting that extracellular cGAMP–STING signaling does not affect neurological development as it pertains to this metric. Consistent with the premature inflammatory signature we observed, 4-month-old *Enpp1*^H362A^ mice exhibited ∼25% reduced rotarod endurance and impaired coordination and fell at ∼25% lower speed than WT. This difference persisted at 22 months of age. At both 4 and 22 months of age, the motor coordination phenotype was rescued in the *Enpp1*^H362A^Sting^gt/gt^ genotype. As an additional assessment of motor coordination requiring postural control and grip, we performed the pole test. Mice were placed head-up at the top of a vertical pole positioned in their home cage, and the time required to turn downward and descend to the base was recorded. Four-month-old *Enpp1*^H362A^ mice exhibited prolonged turn latency compared to WT, which was fully reversed in the *Enpp1*^H362A^Sting^gt/gt^ mice (Fig. 2J). For total descent time, we observed a significant defect when comparing *Enpp1*^H362A^ to *Enpp1*^H362A^Sting^gt/gt^ mice, though this did not reach statistical significance when comparing to WT animals (Fig. 2K, L). In both metrics, the majority of *Enpp1*^H362A^Sting^gt/gt^ animals performed as well as or better than their WT counterparts, consistent with a detrimental effect of STING signaling in general on this test. The defects observed in *Enpp1*^H362A^ mice are consistent with a bradykinesia-like motor phenotype attributed to dysregulated extracellular cGAMP–STING signaling.

Having established motor deficits beginning around 4 months of age in animals lacking extracellular cGAMP hydrolysis by ENPP1, we next assessed the cognitive performance of *Enpp1*^H362A^ mice compared to their WT or *Enpp1*^H362A^*Sting*^gt/gt^ counterparts. We first tested short-term memory in 22-month-old mice using the novel object recognition test, which requires mice to distinguish familiar from novel objects.

Unlike WT mice, all *Enpp1*^H362A^ mice froze when they encountered a novel object (Fig. S2G,H) failing to yield a discrimination index (Fig. S2G). In contrast, *Enpp1*^H362A^*Sting*^gt/gt^ mice behaved similarly to WT mice, demonstrating that unrestrained extracellular cGAMP–STING signaling drives this phenotype. To determine whether *Enpp1*^H362A^ mice failed to explore due to anxiety, locomotor impairment, or a lack of exploration tendency, we next performed the open field test. In contrast to 22-month-old WT mice, *Enpp1*^H362A^ mice rarely explored the center of the open field arena while *Enpp1*^H362A^*Sting*^gt/gt^ exhibited WT-like behavior (Fig. 2M). The *Enpp1*^H362A^ aged mice showed no difference in the distance, speed of their movement, and time spent at walls (Fig. S2I-K), indicating that their deficits in gross locomotor function as measured in the rotarod and pole tests did not contribute to their lack of exploration in this test. These data demonstrate that loss of ENPP1 cGAMP hydrolysis activity causes a phenotype resembling severe anxiety in aged mice via increased extracellular cGAMP-driven STING signaling. This effect was already apparent in 4 month old mice, again consistent with the emergence of the inflammatory signature at this time point (Fig. 2O, S2L-N). Together, these results demonstrate that unregulated extracellular cGAMP–STING signaling initiates behavioral and motor deficits early in life that progressively worsen with age.

### Microglia are one of the major extracellular cGAMP responders via the LRRC8A:C channel

Having established that myeloid cells, especially microglia, are the primary source of extracellular cGAMP in the CNS, we next sought to identify the major cell types receiving the extracellular cGAMP signal. To identify these receivers, we examined which cell types exhibit the most pronounced upregulation of STING-induced genes in response to elevated extracellular cGAMP. To this end, we performed single-cell RNA-seq analysis of the cortex, hippocampus, striatum, and diencephalon from 4- to 6-month-old WT, *Enpp1*^H362A^, and *Enpp1*^H362A^*Sting*^gt/gt^ mice (Fig. 3A). We chose this age to capture near the beginning of the age-related inflammatory and behavioral changes described above in the ENPP1 mutant animals. All major cell types separated into discrete clusters (Fig. 3B-E) identifiable by their molecular markers (Fig. S3A, B). To perform gene expression analysis, we clustered cell types from pooled (Fig. S3C) and individual brain regions (Fig. S3D). While most cell types were present in all of the investigated regions, 85% of inhibitory neurons derive from the striatum (Fig. S3D). Fibroblasts express the highest levels of ENPP1, with many other CNS cell types expressing low to moderate ENPP1 levels (Fig. S3E).

**Figure 3.**
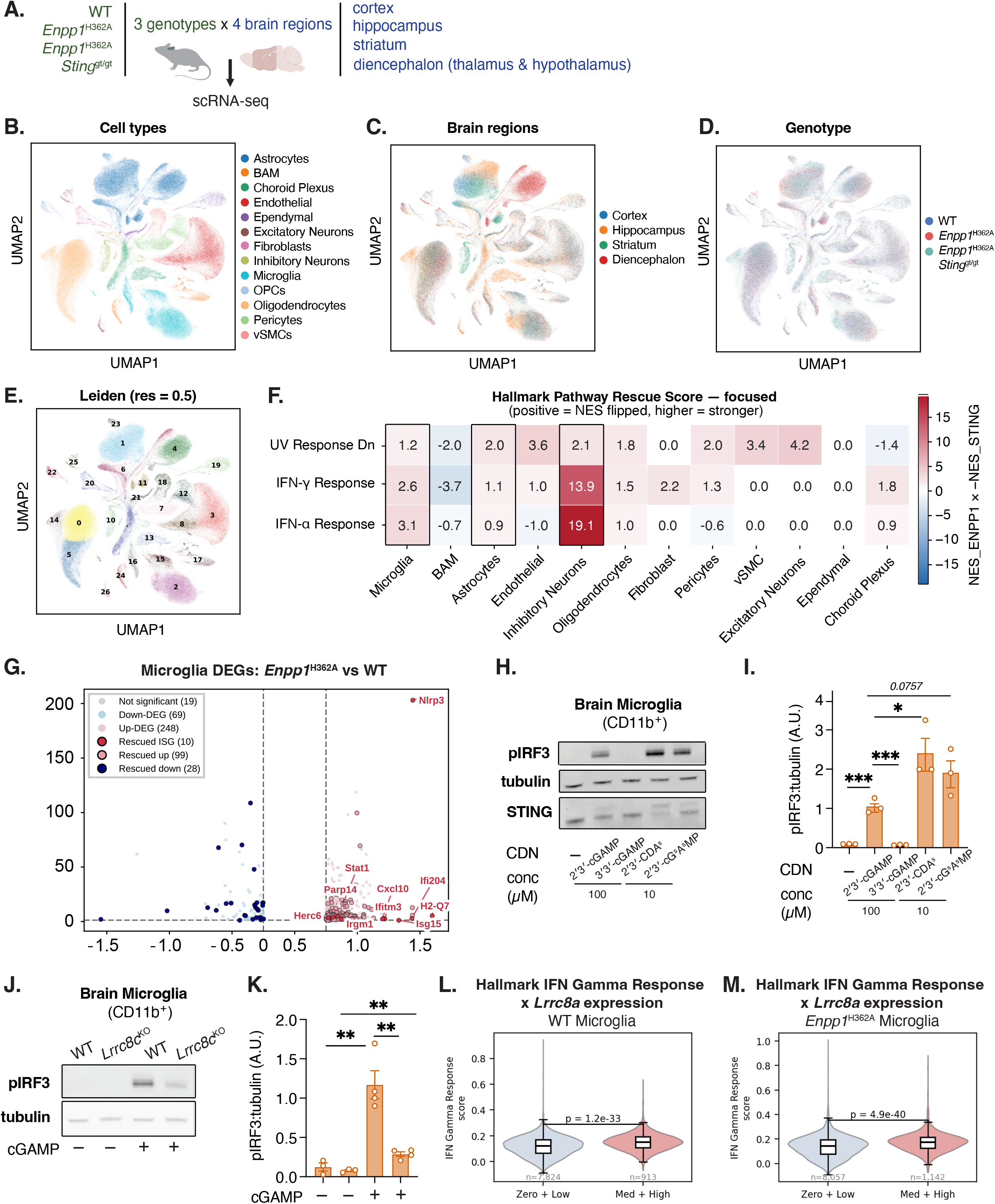
Microglia are one of the major extracellular cGAMP responders via the LRRC8A:C channel. (A) Schematic of scRNA-seq analysis of brain regions of WT, *Enpp1*^H362A^ and *Enpp1*^H362A^ *Sting*^gt/gt^ mice at 4–6 months of age. (B-E) UMAP plots of the annotated clusters of single-cell transcriptomes from wild-type, *Enpp1*^H362A^ and *Enpp1*^H362A^ *Sting*^gt/gt^ mice at 4–6 months of age. Cell types of pooled brain regions of each genotype (B), different brain regions (C), different genotypes (D), and Leiden clusters (E). (F) Hallmark Pathway Rescue Score ( -NES_STING x NES_ENPP1 ) for major cell types of pooled brain regions for each indicated genotype. NES_ENPP1 refers to Hallmark Pathway functional enrichment of *Enpp1*^H362A^ versus wild-type and NES_STING refers to *Enpp1*^H362A^ *Sting*^gt/gt^ versus *Enpp1*^H362A^. (G) Volcano plot showing differentially expressed genes (DEGs) between *Enpp1*^H362A^ versus WTmicroglia at 6 months of age. Genes with FDR values < 0.05 and |log_2_ FC| > 0.25 were called as DEGs for each comparison, and highlighted if rescued by *Enpp1*^H362A^ *Sting*^gt/gt^ . (H,I) Western blot (n=3 biological replicates) for phosphorylated IRF3 (pIRF3), STING, and tubulin as a loading control from MACS-sorted CD11b^+^ primary mouse brain microglia from WT adult mice following incubation with the indicated concentration of cyclic dinucleotides (CDNs). One representative Western blot shown in (H) and quantification of three blots are shown in (I). Data are mean ± s.e.m. *p*-values were calculated using two-sided unpaired t-test with Welch’s correction. (J,K) Western blot (n=2 biological replicates) for pIRF3 and tubulin as loading control from MACS-sorted CD11b^+^ primary mouse brain microglia from WT and *Lrrc8c*^KO^ adult mice following incubation with or without 100 µM 2′3′-cGAMP. One representative Western blot shown in (J) and quantification of multiple blots are shown in (K). Data are mean ± s.e.m. *p*-values were calculated using two-sided unpaired t-test with Welch’s correction. (L,M) Expression of Hallmark IFN-γ response signature in microglia binned by expression of *Lrrc8a*. Hallmark IFN-γ response module score in microglia from wild-type (L) and *Enpp1*^H362A^ (M) mice at 4–6 months of age, all four brain regions pooled. Microglia were binned by log-normalized *Lrrc8a* expression into Zero + Low (≤ 0.5) and Med + High (> 0.5). Inner box plots show the median and interquartile range, with whiskers at 1.5× IQR. P-values were calculated from two-sided Mann–Whitney U tests.

We first took a pathway agnostic approach and investigated all cell types whose gene expression patterns were broadly altered in *Enpp1*^H362A^ mice and restored in *Enpp1*^H362A^*Sting*^gt/gt^ mice as a way to identify extracellular cGAMP responders (Fig. S3F). We observed that oligodendrocytes (3355 genes altered, 60% rescued), astrocytes (2365 genes altered, 46% rescued), endothelial cells (1633 genes altered, 74% rescued), microglia (1072 genes altered, 54% rescued), pericytes (250 genes, 41% rescued) excitatory neurons (128 genes altered, 34% rescued), and inhibitory neurons (103 genes altered, 42% rescued) had the broadest gene expression changes in *Enpp1*^H362A^ mice which were reversed in *Enpp1*^H362A^*Sting*^gt/gt^ mice to different extents, indicating broad, cell type-specific gene expression programs directly or indirectly dependent on extracellular cGAMP-STING signaling.

We then reasoned that the major direct responders to extracellular cGAMP should exhibit higher gene expression of proximal downstream transcriptional programs driven by STING (dsDNA/UV response genes, and IFN-α and IFN-γ signatures) in *Enpp1*^H362A^ mice that are rescued in *Enpp1*^H362A^*Sting*^gt/gt^ mice. We scored cell types that follow this pattern with positive values, where higher responses received larger scores. Among all identified cell types, microglia, astrocytes, inhibitory neurons, and oligodendrocytes exhibited the expected regulatory pattern on these three response signatures (Fig. 3F), identifying them as candidate extracellular cGAMP responders. Interestingly, amongst all cell types, inhibitory neurons have the highest extracellular cGAMP-dependent interferon signature, suggesting that they may be important drivers of IFN production in response to elevated extracellular cGAMP–STING signaling.

Notably, this response resembles the neuronal inflammatory stress response (NISR),^11^ and may help to reconcile previous reports of neuronal cGAS-independent STING signaling associated with neuroinflammation. Due to technical challenges in isolating oligodendrocytes from the mouse brain, we focused on microglia, astrocytes, and neurons as candidate CNS cell types that may directly import extracellular cGAMP and drive downstream STING-dependent responses.

Microglia from 4- to 6-month-old *Enpp1*^H362A^ mice exhibited a stronger STING signature than their WT counterparts, and this signature was abrogated in *Enpp1*^H362A^*Sting*^gt/gt^ (Fig. 3G). Stratifying these cells by brain region, microglia in the striatum showed the strongest increase in the proportion of microglia that were interferon responsive in *Enpp1*^H362A^ mice compared to WT mice and *Enpp1*^H362A^*Sting*^gt/gt^, with the cortex and diencephalon exhibiting milder effects. These results suggest that microglia in the striatum are particularly responsive to elevated extracellular cGAMP, and raise the possibility of region-specific vulnerabilities to neuroinflammatory stimuli (Fig. S3G).

We next sought to determine which transporter is primarily responsible for cGAMP internalization into microglia. To narrow down candidate transporters, we treated primary brain microglia isolated from WT young adult mice (age 2-4 months) with four cyclic dinucleotides (CDNs) known to activate STING – 2′3′-cGAMP, 3′3′-cGAMP, 2′3′-CDA, and 2′3′-cG^s^A^s^MP – and measured their induction of IRF-3 phosphorylation, a major event in STING signaling (Fig. 3H,I). Unlike the 2’-5’-linked CDNs, 3’3’-cGAMP was unable to activate STING signaling outputs, suggesting that it was not taken up by microglia. This substrate fingerprint matched that of the LRRC8A:C channel that we previously identified as the major cGAMP transporter in endothelial cells.^17,22^ Indeed, microglia isolated from *Lrrc8c*^-/-^ mice exhibited only ∼30% of the WT level of pIRF3 in response to extracellular cGAMP supplementation (Fig. 3J,K). Since LRRC8A is the obligatory subunit and LRRC8C requires it for functional channel formation,^30,31^ these results suggest that the LRRC8A:C channel is a major cGAMP importer in isolated microglia.

To probe LRRC8A:C function in cGAMP transport into microglia in vivo, we compared STING signaling in microglia expressing high versus low LRRC8A. LRRC8A-high and LRRC8C-high microglia exhibited significantly elevated IFN-γ signaling in both WT (Fig. 3L, Fig. S3G) and *Enpp1*^H362A^ mice (Fig. 3M, Fig. S3G), and this difference is diminished in *Enpp1*^H362A^*Sting*^gt/gt^ mice (Fig. S3H). Together, these results demonstrate that the LRRC8A:C channel is a major transporter responsible for microglial uptake of and response to extracellular cGAMP. Interestingly, this suggests an intercellular propagation of the cGAMP signal produced primarily by a subset of microglia with elevated cytosolic mtDNA driving cGAS activation and cGAMP production, and internalized by nearby LRRC8A:C-expressing responder microglia to exert its downstream effects.

### Astrocytes can import extracellular cGAMP through system xc-facilitated by glutamate transporters

Astrocytes have been implicated in STING-mediated neuroinflammation, including through astrocyte-intrinsic cGAS–STING signaling.^32^ However, our observation that astrocytes upregulate the STING signature in the absence of extracellular ENPP1 cGAMP hydrolysis activity strongly suggests that extracellular cGAMP also influences astrocytes. Astrocytes from 4- to 6-month-old *Enpp1*^H362A^ mice exhibited expression changes in nearly a thousand genes compared with WT mice and these changes were largely rescued in *Enpp1*^H362A^*Sting*^gt/gt^ mice (Fig. 4A). In particular, these differentially expressed genes included upregulation of ISGs and genes participating in the glutamate–glutamine cycle (Fig. 4B). In agreement with our scRNAseq data, primary mouse astrocytes responded to extracellular cGAMP treatment, as measured by the induction of pIRF3 (Fig. 4C). We first asked whether the LRRC8A:C channel responsible for cGAMP import in microglia also mediates cGAMP transport in astrocytes.

**Figure 4.**
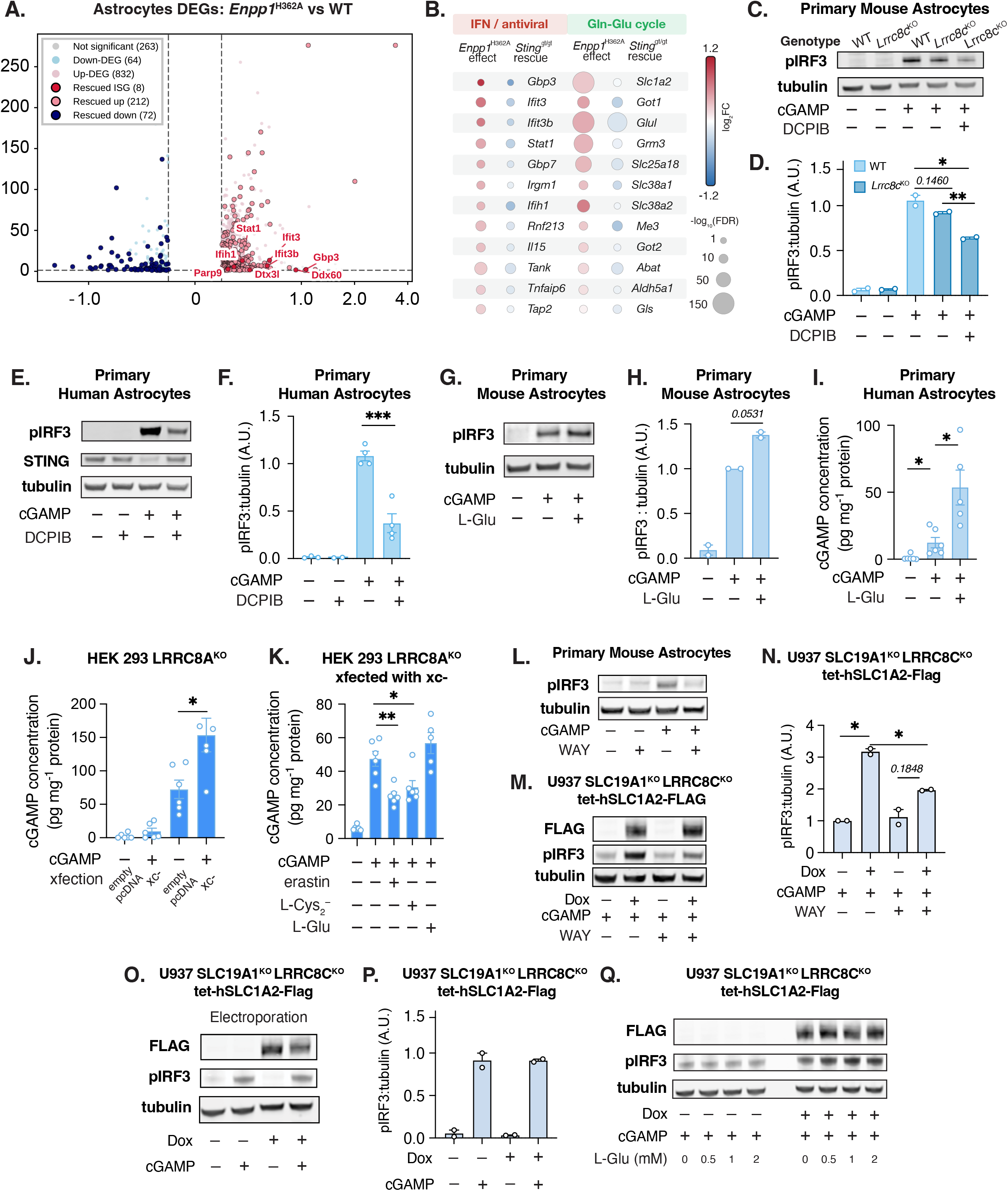
Astrocytes can import extracellular cGAMP through system xc-facilitated by glutamate transporters. (A) Volcano plot showing *Enpp1* effect (*Enpp1*^H362A^ versus wild-type) DEGs in astrocytes at 4–6 months of age. Genes with FDR values < 0.05 and |log_2_ FC| > 0.25 were called as differentially expressed genes for each comparison. (B) Bubble plot showing STING-dependent DEGs in astrocytes by comparing *Enpp1* effect (*Enpp1*^H362A^ versus wild-type) and *Sting*^gt/gt^ effect (*Enpp1*^H362A^ *Sting*^gt^ versus *Enpp1*^H362A^). (C,D) STING signaling output as measured by pIRF3 levels in MACS-sorted ACSA-2^+^ primary mouse brain astrocytes from WT and *Lrrc8c*^KO^ adult mice with or without treatment with 100 µM 2′3′-cGAMP and/or 20 µM DCPIB. One representative Western blot shown in (C) and quantification of n = 2 biological replicates is shown in (D). (E,F) STING signaling output as measured by pIRF3 levels in primary human brain astrocytes with or without treatment with 100 µM 2′3′-cGAMP and/or 20 µM DCPIB One representative Western blot is shown in (E) and quantification of n = 4 biological replicates is shown in (F). (G,H) STING signaling output as measured by pIRF3 levels in MACS-sorted ACSA-2^+^ primary mouse brain astrocytes from WT adult mice with or without treatment with 100 µM 2′3′-cGAMP and/or 1-2 mM L-glutamate. One representative Western blot is shown in (G) and quantification of n = 2 biological replicates is shown in (H). (I) Intracellular 2′3′-cGAMP concentrations in human primary astrocytes after treatment with or without 10 µM 2′3′-cGAMP and/or 1-2 mM L-glutamate, measured by enzyme-linked immunosorbent assay (ELISA) (n = 5–6 biological replicates). (J) Intracellular 2′3′-cGAMP concentrations in HEK 293 LRRC8A^KO^ cells expressing system xc- (SLC7A11 and SLC3A2) after treatment with 10 µM 2′3′-cGAMP, measured by enzyme-linked immunosorbent assay (ELISA) (n = 5–6 biological replicates). (K) Intracellular 2′3′-cGAMP concentrations in HEK 293 LRRC8A^KO^ cells expressing system xc-(SLC7A11 and SLC3A2) after treatment with or without 10 µM 2′3′-cGAMP and/or erastin (50 µM) and/or L-cystine (1 mM) and/or L-glutamate (1-2 mM), measured by enzyme-linked immunosorbent assay (ELISA) (n = 5–6 biological replicates). (L) STING signaling output as measured by pIRF3 levels in MACS-sorted ACSA-2^+^ primary mouse brain astrocytes from WT adult mice treated with or without 100 µM 2′3′-cGAMP and/or 20 µM WAY-213613. One representative Western blot is shown out of n = 2 biological replicates. (M,N) STING signaling output as measured by pIRF3 levels in U937 *SLC19A1*^KO^ *LRRC8A*^KO^ tet-h*SLC1A2*-Flag cells treated with or without 2 µg/ml doxycycline (Dox) for 72 h, and treated with or without 50 µM 2′3′-cGAMP and/or 1 µM WAY-213613 for 1h. . One representative Western blot is shown in (M) and quantification of n = 2 biological replicates is shown in (N). (O,P) STING signaling output as measured by pIRF3 levels in U937 *SLC19A1*^KO^ *LRRC8A*^KO^ tet-h*SLC1A2*-Flag cells treated with or without 2 µg/ml doxycycline (Dox) for 72 h, and electroporated with 100 nM 2’3’-cGAMP.One representative Western Blot is shown in (O), and quantification of n = 2 biological replicates is shown in (P). (Q) STING signaling output as measured by pIRF3 levels in U937 *SLC19A1*^KO^ *LRRC8A*^KO^ tet-h*SLC1A2*-Flag cells treated with or without 2 µg/ml doxycycline (Dox) for 72 h, and treated with 50 µM 2′3′-cGAMP and/or the indicated concentrations of L-glutamate.(n = 2 biological replicates). One representative Western blot out of n = 2 biological replicates is shown. Data are mean ± s.e.m. p-values were calculated using two-sided unpaired Student’s t-tests unless otherwise indicated.

Astrocytes from *Lrrc8c*^-/-^ mice did not exhibit a decrease in response to extracellular cGAMP. Since the LRRC8C subunit is required to form a complex with the LRRC8A subunit to transport cGAMP, our results indicate that LRRC8A:C is not a major cGAMP transporter in astrocytes. However, treatment with a non-specific LRRC8A:C inhibitor, DCPIB, markedly reduced the ability of extracellular cGAMP to elicit STING signaling in both human and mouse primary astrocytes, suggesting that a different molecular target of DCPIB may contribute to cGAMP import (Fig. 4C-F).

DCPIB was previously shown to nonspecifically inhibit other glutamate transporters including SLC1A2.^33^ We reasoned that the addition of excess extracellular glutamate could help determine whether one or more glutamate transporters (EAATs) might be involved in cGAMP import by providing a cis-inhibitory effect.^34^ Surprisingly, extracellular glutamate boosted cGAMP-mediated p-IRF3 signaling in primary mouse astrocytes (Fig. 4G, H) as well as directly increased cGAMP transport in primary human astrocytes (Fig 4I) – the opposite of what we would expect for cis-inhibition of a glutamate transporter that can also transport cGAMP. This result led us to consider the possibility that glutamate import via SLC1A2 or another glutamate importer may facilitate cGAMP import by feeding a cGAMP-glutamate antiporter in trans. System xc^−^ is the only known glutamate exchanger, which has been demonstrated to export one glutamate coupled to the import of one cystine molecule.^35^

To establish whether system xc-can indeed transport cGAMP, we turned to HEK293 cells as a model system: cGAMP import into these cells relies partially on endogenous LRRC8A:C channels,^17^ so we used a knockout line with transient overexpression of both SLC7A11 and SLC3A2 (constituting system xc-) or empty pcDNA as control. Upon SLC7A11 and SLC3A2 dual overexpression, cGAMP uptake was increased compared to the empty DNA control (Fig. 4J) and this increase was inhibited by the specific system xc-inhibitor erastin,^36^ inhibited by SLC7A11’s canonical substrate cystine, and boosted by glutamate (Fig. 4K). Similarly, overexpression of both SLC7A11 and SLC3A2, but less so SLC7A11 or SLC3A2 alone, in these cells also increased cGAMP-mediated p-IRF3 signaling (Fig. S4A).

We wondered if SLC1A2, which is known to be inhibited by DCPIB and is the most abundant glutamate transporter in astrocytes, facilitates cGAMP transport by system xc^−^ in trans; if true, this may explain the counterintuitive DCPIB result above. Indeed, WAY-213613, a specific SLC1A2 inhibitor,^37^ abolished the induction of pIRF3 in mouse primary astrocytes (Fig. 4L). Based on these findings, we hypothesized that any cell line with system xc^−^ expression could achieve enhanced cGAMP import activity if they overexpressed SLC1A2. We generated a doxycycline-inducible SLC1A2 overexpression system in a U937 model cell line carrying knockouts of its two dominant cGAMP transporters, SLC19A1 and LRRC8A:C^17,20^ (U937 SLC19A1^-/-^ LRRC8C^-/-^ tet-SLC1A2-Flag). Indeed, doxycycline induction of SLC1A2 expression led to an increase of phosphorylated IRF3, which was blocked by WAY-213613 (Fig. 4M,N) as well as erastin (Fig. S4B). This enhancement was abolished when cGAMP was electroporated directly into cells, bypassing transporter requirements (Fig. 4O,P), suggesting that SLC1A2 facilitates cGAMP internalization instead of its downstream signaling. Indeed, addition of extracellular glutamate further boosted the effect of SLC1A2 overexpression (Fig. 4Q) as seen in primary astrocytes. Together with the ability of DCPIB and WAY-213613 to inhibit extracellular cGAMP signaling in astrocytes, this data suggests an important role for SLC1A2 as a facilitator that supports system xc- to uptake extracellular cGAMP by providing the necessary intracellular glutamate concentration to drive antiport.

### Extracellular cGAMP can activate neuronal STING through system xc^_^ facilitated by glutamine transporters and glutaminase

Neurons can produce the NISR associated with neuroinflammation, with interferons constituting the main signaling axis in this response.^11^ Previous work showed that neuronal STING is important in the pathology of EAE/MS, but it was hypothesized to act through cGAMP-independent activation pathways.^11^ Our scRNAseq data offers another possibility: that neurons might be directly responding to elevated extracellular cGAMP in the contexts of EAE/MS and aging to induce a NISR-like gene expression program that promotes neuroinflammatory phenotypes.

To test whether extracellular cGAMP can directly activate neuronal STING, we treated mouse primary neurons isolated from P1 cortex and striatum with low concentrations of extracellular cGAMP plus the ENPP1 inhibitor STF-1623 to limit cGAMP degradation. Fifteen minutes after treating neurons with cGAMP, we detected intracellular cGAMP, which was below the detection limit in untreated neurons, suggesting that neurons may directly import extracellular cGAMP (Fig. 5A). Twenty four hours after treatment, we observed hundreds of differentially expressed transcripts with ≥2-fold changes; the most upregulated, including *Ifit44*, *Irf7*, and *Ifit3*, which were induced >10-fold, belong to the IFN-α, IFN-γ, and downstream STING signaling pathways, suggesting neurons may respond to cGAMP (Fig. 5B, Fig. S5A).

**Figure 5.**
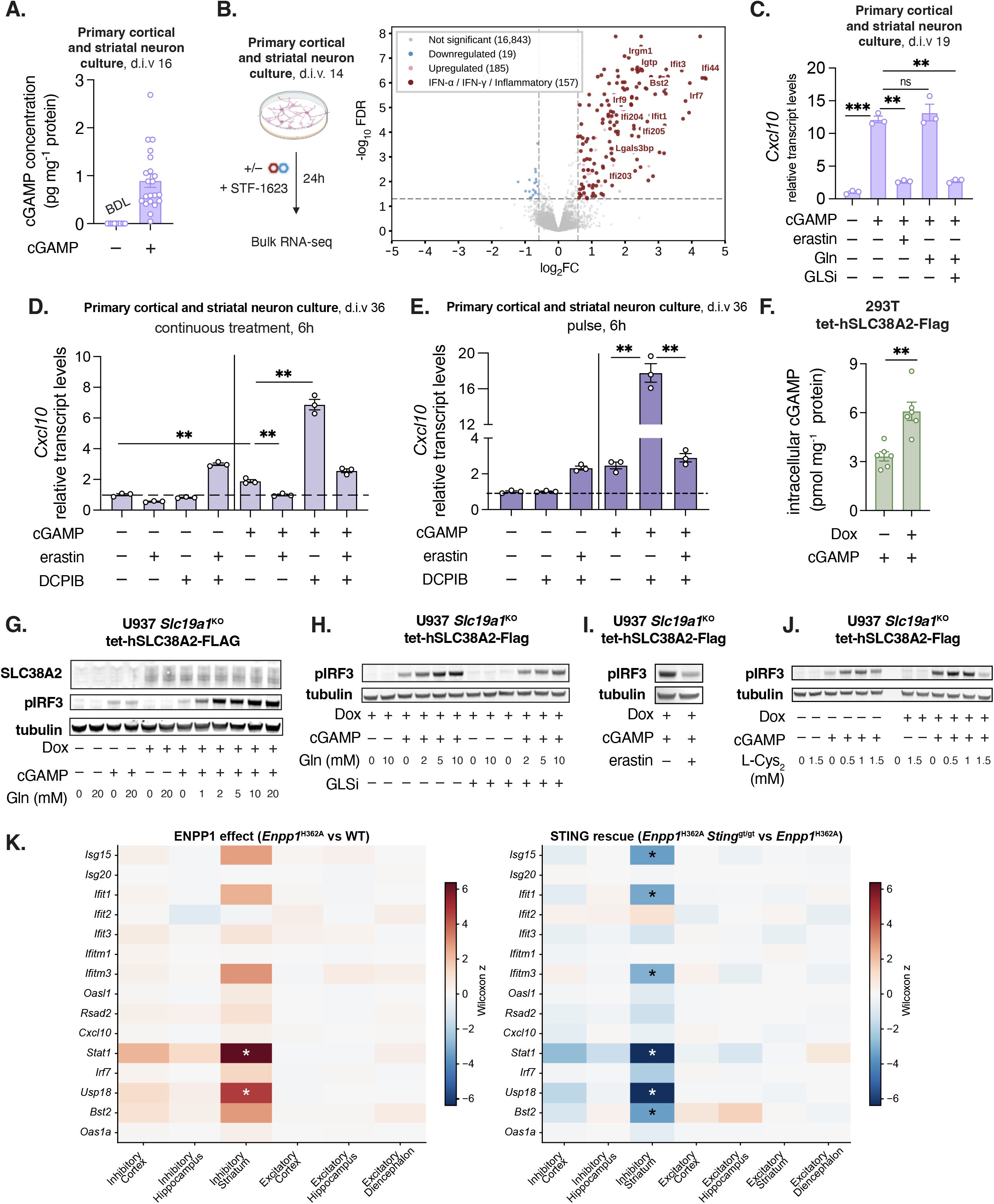
Extracellular cGAMP can activate neuronal STING through system xc^_^ facilitated by glutamine transporters and glutaminase. (A) Quantification of intracellular 2′3′-cGAMP concentrations in primary mouse cortical and striatal neurons following 15-minute treatment with 10 µM 2′3′-cGAMP by enzyme-linked immunosorbent assay (ELISA) and normalized to total protein amount in the cell lysates (n=21 biological replicates). (B) Bulk RNA sequencing volcano plot of primary mouse cortical and striatal neuron cultures after 24 h treatment with 10 µM 2′3′-cGAMP and 1 µM ENPP1 inhibitor STF-1623. Genes with FDR values < 0.05 and |log_2_ FC| > 1 were considered differentially expressed after 2′3′-cGAMP treatment. (C) *Cxcl10* transcript levels determined by RT-qPCR of primary mouse cortical and striatal neuron cultures after pre-treatment with glutaminase inhibitor telaglenastat (CB-839, 10µM), followed by L-Gln (2 mM) or erastin (50 µM) for 15 min, then 2′3′-cGAMP for 6h (n=3 technical replicates per condition, replicated across 3 independent experiments). (D) *Cxcl10* transcript levels determined by RT-qPCR of primary mouse cortical and striatal neuron cultures after pre-treatment with erastin (50 µM), DCPIB (20 µM), or both for 15 min, followed by 2′3′-cGAMP for 6h (n=3 technical replicates per condition). (E) *Cxcl10* transcript levels determined by RT-qPCR of primary mouse cortical and striatal neuron cultures after pre-treatment with erastin (50 µM), DCPIB (20 µM), or both for 15 min, followed by 2′3′-cGAMP for 30 min. All compounds were washed out and cells were incubated in compound-free media for 5h (n=3 technical replicates per condition). (F) Intracellular 2′3′-cGAMP concentration in 293T tet-h*SLC38A2*-Flag cells as measured by ELISA following incubation with or without 2 µg/ml doxycycline (Dox) for 18 h and treatment with or without 10 µM 2′3′-cGAMP for 15 min in the presence of ENPP1 inhibitor STF-1623. Measured values are normalized to total protein amount in the cell lysates (n = 6 biological replicates). (G) Western blot assessing STING signaling through pIRF3 levels and SLC38A2 expression in U937 *Slc19a1*^KO^ tet-h*SLC38A2*-Flag cells following incubation with or without 2 µg/ml doxycycline (Dox), the indicated concentrations of L-Glutamine, and 50 µM 2′3′-cGAMP . One representative Western blot of n = 2 biological replicates is shown. (H) Western blot assessing STING signaling through pIRF3 levels in U937 *Slc19a1*^KO^ tet-h*SLC38A2*-Flag cells following incubation with or without 2 µg/ml Dox, 50 µM 2′3′-cGAMP, the indicated concentrations of L-Glutamine, and the glutaminase inhibitor telaglenastat (GLSi). One representative Western blot of n = 2 biological replicates is shown. (I) Western blot assessing pIRF3 levels in U937 *Slc19a1*^KO^ tet-h*SLC38A2*-Flag cells following incubation with 2 µg/ml doxycycline (Dox) and cGAMP, with or without erastin (50µM). One representative Western blot of n = 2 biological replicates is shown. (J) Western blot to measure pIRF3 in U937 *Slc19a1*^KO^ tet-h*SLC38A2*-Flag cells following incubation with or without Dox and cGAMP, plus the indicated concentrations of L-cystine. One representative Western blot of n = 2 biological replicates is shown. (K) Heatmap of the Wilcoxon rank-sum z-score for a 15-gene ISG panel; all panel genes are shown regardless of expression level. Left, *Enpp1*^H362A^ vs. WT; right, *Enpp1*^H362A^ *Sting*^gt/gt^ vs. *Enpp1*^H362A^. Positive (red) z indicates higher expression in the first-named genotype. *: FDR < 0.05, Benjamini–Hochberg correction across all genes in each comparison. Region × neuron-type combinations with fewer than 30 cells per genotype were not tested and are not shown. Data are mean ± s.e.m. *p*-values were calculated using two-sided unpaired Student’s t-tests unless otherwise indicated.

To further test whether neurons can take up cGAMP through regulated mechanisms, we sought to identify a neuronal cGAMP transporter. We first investigated whether any known cGAMP transporters are expressed in neurons. Our bulk RNAseq data shows that these primary mouse neurons robustly express the LRRC8A:C channel, as well as the system xc-antiporter we showed above to be a major contributor to microglia and astrocyte cGAMP import, respectively. We hypothesized that the high cytosolic glutamate concentrations in neurons (∼10 mM, with higher concentrations in synaptic vesicles) could facilitate system xc-’s cGAMP import in exchange for glutamate export. Indeed, the SLC7A11 inhibitor erastin reduced extracellular cGAMP-mediated ISG expression in primary cortical mouse neurons by more than 50% (Fig. 5C), suggesting that system xc- is also an important cGAMP importer in neurons.

The high intracellular concentration of glutamate in neurons is generated through conversion of glutamine to glutamate by the cytosolic enzyme glutaminase. The glutaminase inhibitor telaglenastat completely inhibited the ability of primary mouse cortical neurons to induce the ISG transcript *Cxcl10* in response to extracellular cGAMP (Fig. 5C), supporting the idea that glutamate-cGAMP antiport is a primary means by which neurons import extracellular cGAMP. Addition of excess glutamine did not boost extracellular cGAMP signaling, which we attribute to either the high glutamine concentration already present in the culture media or the high intracellular glutamine in neurons at homeostasis. In addition, the (non-specific) LRRC8A:C channel inhibitor DCPIB boosted cGAMP induced ISG expression (Fig. 5D), suggesting that LRRC8A:C or another DCPIB target could be a cGAMP exporter in neurons instead of an importer. To avoid any indirect effects of erastin and DCPIB outside cGAMP transport, we provided a 30-min pulse of cGAMP together with erastin or DCPIB or both, and assayed *Cxcl10* expression in the neurons 6 hours later. Strikingly, DCPIB greatly boosted cGAMP-induced ISG expression while erastin abolished all the effects of DCPIB (Fig. 5E), suggesting they act directly on cGAMP transport instead of its downstream signaling which occurs after 30 min.^38^ Primary neuronal cultures supplemented with cytosine arabinoside (AraC) to inhibit glial proliferation contain low levels of astrocytes (∼5–10%). While we cannot exclude a minor contribution of astrocytes to these data, our findings with glutaminase inhibition, together with the opposing effect of DCPIB in neurons versus astrocytes, argue against astrocytes being the major contributors to what we observed. Together, our results suggest that the system xc-antiporter imports cGAMP into neurons in exchange for glutamate export while the LRRC8A:C channel and/or other DCPIB targets may act to clear intracellular cGAMP from neurons.

Neurons highly express SLC38A1 (SNAT1) and SLC38A2 (SNAT2) and use them to import extracellular glutamine, feeding the cytosolic production of glutamate for neurotransmission. We sought to overcome the confounding effect of glutamine concentration in primary neuronal culture and formally test whether glutamine import through SLC38A1/2 facilitates system xc- -mediated cGAMP import. To do so, we overexpressed SLC38A2 under a doxycycline-inducible promoter in 293T or U937 cells lacking their endogenous cGAMP transporter, SLC19A1. Notably, both of these cell types express system xc- and are also known to express high levels of glutaminase.^39^ Inducing SLC38A2 greatly increased intracellular cGAMP levels (Fig. 5F). It also increased IRF3 phosphorylation in a glutamine dependent manner: higher concentrations of glutamine increased the signal while SLC38A2 overexpression did not have any effect without extracellular glutamine (Fig. 5G). Like in primary neurons, the glutaminase inhibitor telaglenastat completely abolished SLC38A2’s effect on cGAMP signaling (Fig. 5H). Finally, both erastin (Fig. 5I) and cystine (Fig. 5J) inhibited SLC38A2-aided cGAMP signaling in these cells, suggesting that SLC38A2 boosts cGAMP signaling through system xc-instead of directly transporting cGAMP. Together, these results illustrate how extracellular cGAMP utilizes the glutamine–glutamate cycle to enter both neurons and astrocytes to elicit non-cell-autonomous inflammatory responses in the CNS.

The mouse primary cortical neuron population includes both inhibitory and excitatory neurons, but our extracellular cGAMP-dependent pathway analysis above (Fig 3F) clearly showed that the *in vivo* neuronal response can be primarily attributed to the inhibitory neuron class. We next asked whether the inhibitory neurons across different brain regions show differential responses to elevated extracellular cGAMP. Compared to other regions, striatal inhibitory neurons showed the strongest increase in ISG expression in *Enpp1*^H362A^ mice compared to WT, which was reversed in *Enpp1*^H362A^*Sting*^gt/gt^ (Fig. 5K, S5B). Two non-exclusive explanations for this regional vulnerability are possible. First, the striatum contains the highest density of inhibitory neurons in the brain, which may permit greater paracrine amplification of interferon signaling among neighboring responders. Second, the striatum contains the largest proportion of interferon-responsive microglia in response to extracellular cGAMP elevation, which we expect would contribute to a more inflammatory microenvironment in the striatum that could promote further signal amplification throughout the tissue (Fig. S3C). Notably, an outsized striatal vulnerability to elevated extracellular cGAMP could help explain the phenotypes we observed in the pole test^40^ and rotarod test^41^ for *Enpp1*^H362A^ animals. In addition, because striatal inflammation and synaptic injury occur in EAE,^42^ and striatal microglia^43^ and astrocytes^44^ have been reported to acquire inflammatory features with age, this response may represent a shared contribution to EAE- and aging-associated neuropathology.

## DISCUSSION

In this work, we established extracellular cGAMP as a central immunotransmitter in the neuroinflammation that contributes to neurodegenerative disease and brain aging. We show that microglia and other encephalitogenic myeloid cells produce and export large quantities of cGAMP in response to mitochondrial DNA leakage. Accumulated extracellular cGAMP can then activate STING signaling in other microglia, astrocytes, and inhibitory neurons via cell type-specific transporters that, in astrocytes and neurons, appear to utilize the glutamine-glutamate neurotransmission cycle to facilitate cGAMP transport.

Separate from its canonical role as an intracellular second messenger, the strictly compartmentalized extracellular cGAMP paracrine signaling axis is specifically under the control of the extracellular enzyme ENPP1 in the CNS. We use this selective control of the extracellular cGAMP pool as a lever to pinpoint its functional role independent of intracellular cGAS–STING signaling. Interestingly, cGAMP lacks intracellular hydrolases, highlighting the evolutionary importance of restraining its accumulation in the extracellular space to protect against metabolic tissue dysfunction,^45^ neuroinflammation, and potentially other deleterious consequences yet to be discovered. While high ENPP1 activity can promote tumor immune evasion and immunotherapy resistance through attenuating the innate immune cascade,^13–15,46^ here we show that low or absent ENPP1 activity can accelerate age- and disease-related inflammatory processes. The stronger responses that we observed in the striatum compared to other brain regions suggests that additional neuroinflammatory disorders with clear striatal pathology, such as Huntington’s^47^ and Parkinson’s^48^ may be candidate contexts in which to explore the role of extracellular cGAMP in future work. We hypothesize that physiological levels of ENPP1^14^ and its paralog ENPP3^24^ act as an innate immune checkpoint throughout the body to maintain low levels of extracellular cGAMP, striking a fine balance to maintain paracrine immune communication while restraining inflammatory signaling cascades at baseline. An inverse association has been reported between certain cancers and neurodegenerative diseases, particularly Alzheimer’s disease.^49^ Although its basis remains unresolved, this epidemiologic pattern raises the possibility that opposing deviations in extracellular cGAMP signaling could contribute to cancer susceptibility and neuroinflammatory disease.

While in this study we focused on CNS-specific cell types as receivers of the extracellular cGAMP signal, it is possible that other nearby cell types may also contribute. Our previous work identified endothelial cells as prominent responders to ENPP1-regulated extracellular cGAMP signaling in breast tumors, exhibiting a strong interferon-stimulated gene response.^14^ It remains to be determined whether endothelial cells in the CNS act in the same manner. Perhaps counterintuitively, we identified microglia as both the primary cGAMP producing cells and as key responders to extracellular cGAMP. The effect of losing extracellular cGAMP hydrolysis activity through the ENPP1^H362A^ genotype shows that, although intracellular cGAMP-STING signaling surely does occur within cGAMP-producing microglia, a portion of the cGAMP signal also travels to nearby microglia to activate STING non-cell-autonomously – and this extracellular portion is functionally important for the phenotypes we observed. We postulate that cGAMP export may be thermodynamically favored by the electrochemical gradient of cGAMP, and future work will be required to pinpoint any determinants of cGAMP export in this context, as well as how far and through which means the soluble cGAMP signal can spread within (and potentially beyond) the CNS.

We previously identified the LRRC8A:C channel as the cGAMP transporter in endothelial cells.^17^ Here, we showed that the LRRC8A:C channel is also responsible for importing cGAMP in microglia. Our evidence in neurons that the non-specific LRRC8A:C inhibitor DCPIB significantly increased – rather than decreased – cellular responses to extracellular cGAMP also raises the possibility that this same channel may function bidirectionally to export cGAMP when intracellular concentrations are high; future work will be required to investigate this further. Consistent with our prior studies that highlighted cell type-specific cGAMP import, we identify system xc- as a cGAMP/glutamate antiporter that controls cGAMP uptake in astrocytes and neurons. We further uncover distinct cell-type-specific metabolic circuits that supply the intracellular glutamate required to drive this transport process. Together, these data establish that astrocytic glutamate uptake and neuronal glutamine uptake fuel a shared system xc- -dependent cGAMP import pathway through distinct metabolic inputs. These findings raise the possibility that astrocytes and neurons may tune their cGAMP responsiveness through different nutrient-entry pathways: by local extracellular glutamate availability in astrocytes, but by glutamine access and glutaminase activity in neurons.

Our discovery of the role of extracellular cGAMP in neuroinflammation underscores three potential nodes of regulation that may be amenable to therapeutic intervention. First, we show a pronounced delay of EAE-related paralysis phenotypes when treating mice with a cGAS inhibitor, demonstrating a therapeutic effect of lower cGAMP production even after the inflammatory cascade has been initiated. Two cGAS inhibitors have exhibited strong safety profiles in phase I and are currently in phase II clinical trials for treating autoimmune syndromes.^50–52^ However, these existing inhibitors do not cross the blood-brain barrier; our study urges the development of brain-penetrating cGAS inhibitors to explore their potential therapeutic activity in neuroinflammatory contexts. Second, we showed a striking reversal of extracellular cGAMP-mediated neuroinflammation phenotypes when paired with the inactive STING allele, providing strong genetic evidence that inhibiting STING directly may resolve neuroinflammation. Despite being a major focus of many pharmaceutical companies, no STING inhibitors have yet advanced into clinical trials, suggesting that more mechanistic studies are required to aid in STING inhibitor drug discovery.

Finally, provision of increased cGAMP hydrolase activity in the brain through upregulation of endogenous ENPP1, local delivery of a cGAMP hydrolase, or infiltration of cell therapies engineered to express high levels of ENPP1 may be another practical strategy to quell neuroinflammation.

This study establishes for the first time a functional role for the extracellular cGAMP axis in the central nervous system, defining the primary flow of inflammatory information from microglia to other microglia, astrocytes, and neurons. Through this work, we begin to illuminate the tissue-wide orchestration of neuroinflammatory responses in aging and disease, which we hope will inspire further exploration of this mechanism, the other neuroinflammatory contexts it may influence, and therapeutic opportunities that harness these findings to tackle neuroinflammation from a new angle.

### Limitations of the study

We have not tested the significance of the identified transporters in vivo nor how endogenous cystine, glutamine, and glutamate affect cGAMP transport in vivo and vice versa. This is because cystine, glutamine, and glutamate are vital brain metabolites with tightly regulated concentrations in the brain, making it difficult to perturb their transporters genetically or concentrations pharmacologically. Future studies developing cGAMP-selective mutations of SLC7A11 are warranted to address these questions.

## RESOURCE AVAILABILITY

### Materials availability

• ● Plasmids generated in this study will be made available upon request.

### Data and code availability

● This paper analyzes existing, publicly available data, accessible at https://www.proteinatlas.org/ and https://doi.org/10.5281/zenodo.8338963.
● All data reported in this paper will be shared by the lead contact upon request.

## Supporting information

Table S4. Curated gene sets

## ACKNOWLEDGMENTS

We thank Thaiss, Konermann, Li Lab members, C. Ricci-Tam, B. Plosky, and the Scientific Publication Team at Arc Institute for feedback throughout the course of this study. We thank our funding support, including Arc Institute (D.R.C., R.W., N.L., Y.L., A.S., A.M., S.S., H.G., C.T., S.K., L.L.), Stanford Graduate Fellowship (D.R.C.), and Sarafan ChEM-H Chemistry/Biology Interface Training Program (D.R.C). We thank M. Gilbert at the Arc FACS facility. We thank C. Soekler for support with behavioral assay setups. We thank G. Skariah and G. Grenot for colony management and assistance with disease models and pharmacological treatments. We thank R. E. Mardjuki, J. A. Carozza, and T. Lan for synthesizing cGAMP. We thank X. Cao for thoughtful insight on characterization of transporters.

## AUTHOR CONTRIBUTIONS

Conceptualization, D.R.C., R.W., and L.L.; methodology, D.R.C., R.W., N.L., A.S., S.S., S.K., and L.L.; Investigation, D.R.C., R.W., N.L., Y.L., T.Y., A.S., A.M., S.S. A.P., S.K., and L.L.; writing—original draft, D.R.C., and L.L.; writing—review & editing, D.R.C., A.P., L.L.; funding acquisition, D.R.C., H.G., C.T., S.K., L.L.; resources, H.G., C.T., S.K., L.L.; supervision, H.G., C.T., S.K., L.L.

## DECLARATION OF GENERATIVE AI AND AI-ASSISTED TECHNOLOGIES

The authors did not use AI in preparation of the manuscript.

## METHODS

### Mouse models

C57BL/6J (Stock #000664) and C57BL/6J-Sting1gt/J (Stock #017537) mice were purchased from the Jackson Laboratory. Lrrc8c^KO^ (B6.129S2-Fad158<tm1Maim>/7Maim, RBRC05143) was purchased from Riken. *Enpp1*^H362A^ mice were previously generated^13^ and characterized in-house and bred with C57BL/6J-Sting1gt/J to generate *Enpp1*^H362A^ x *Sting*^gt/gt^. Only female mice were used for *in vivo* studies, and both male and female mice were used for primary cell isolation where appropriate. Mice were maintained at Stanford University and Arc Institute in compliance with the Stanford University and Arc Institute Institutional Animal Care and Use Committees’ regulations. All procedures were approved by the Stanford University and Arc Institute Administrative Panels on Laboratory Animal Care.

### Experimental Autoimmune Encephalomyelitis (EAE)

Female C57BL/6 mice (wild-type, *Enpp1*^H362A^ and *Enpp1*^H362A^ x *Sting*^gt/gt^) aged 9–11 weeks were used for EAE induction. Mice were immunized with Hooke kits (an emulsion of MOG_35–55_ in CFA and Pertussis Toxin in PBS, Hooke Laboratories #EK-2110) according to the manufacturer’s instructions. Briefly, 0.2 ml of the MOG_35–55_ in CFA emulsion was injected subcutaneously at two different sites, followed by intraperitoneal injection of 100 ng pertussis toxin. Another 100 ng pertussis toxin was intraperitoneally injected 48h later. Mice were scored daily, starting on day 11 post-immunization.

For symptomatic treatment, wild-type female C57BL/6 mice were treated intraperitoneally with RU.521 (Invivogen, #inh-ru521-2), dissolved in 10% DMSO, 40% PEG_300_, 5% Tween-80, and 45% Saline) at 10 mg/kg or vehicle on day 11 after EAE induction, when the mice exhibited symptoms defined as a clinical score between 0 and 1. Mice were randomly assigned to different experimental treatments and scored by an investigator blinded to their treatment or genotype. All treatments were carried out on at least four mice and in at least two independent experiments, as indicated in the figure captions.

Clinical scores were defined as follows: 1 = limp tail; 2 = hind limb weakness (inability to fully spread apart hindlimbs); 3 = complete paralysis of at least one of the hindlimbs or partial paralysis of both hindlimbs with inability to maintain a straight gait; 4 = complete hind limb paralysis; and 5 = found dead or moribund and/or a score of 4 for two consecutive days. Easy access to wet food and water was provided for the animals throughout disease progression.

### Open Field Test

During the light phase, mice were moved into and acclimatized to a dimly lit testing room for 30 min being placed into a 40 × 40 cm box with white opaque walls. Mice were allowed to explore freely for 10 min while the investigator left the testing room. Tracking and analysis were conducted using the EthoVision XT software (Noldus).

### Weight measurements

Aged mice did not show any significant difference in their weight distribution across the three genotypes tested in this study. Only mice that weighed between 25–40g were subjected to rotarod and pole tests.

### Accelerating rotarod

Rotarod testing occurred in two consecutive days, one training day and one formal test day, where motor evaluation and scoring was performed. During the light phase, mice were acclimatized to the room for 1 h before going on the rotarod. Rotarod performance was tested using a five-station rotarod treadmill. During training day 1, mice were allowed to acclimate to the rotarod for 5 min at a constant speed of 4 rpm. If a mouse fell, it was placed back on the rod up until the 5 min of training concluded. On the day of evaluation, the rod accelerated from 4 to 40 rpm at a constant rate of acceleration over 300 s. Testing consisted of three trials per day, separated by 2 min each. The rotarod was sanitized in between each trial. A trial was terminated when a mouse fell off, made one complete backward revolution while hanging on, or after 300 s (maximum speed, no further acceleration).

### Pole test

A vertical metal pole (1 cm diameter, 50–60 cm height) was mounted on a base sized to fit within a standard mouse cage and wrapped along its full length with an adhesive non-slip liner to provide grip; the liner was secured at both ends with tape and replaced as it wore. Testing was performed in the home cage, with bedding pushed aside and the pole positioned upright in the center. The pole and base were wiped with 70% ethanol and re-lined between each mouse tested. Pole test occurred in two consecutive days, one training day and one formal test day, where motor evaluation and scoring was performed. On each training and test day, mice were transferred from the vivarium to the behavior room at least 1 h before testing to acclimate, and individual mice were allowed to explore the cage containing the apparatus for 1 min prior to each session. On training day 1, each mouse underwent five trials in which it was placed head-down on the pole at progressively higher positions (10, 20, 30, and 40 cm) and then at the top (three times), and allowed to descend freely; the experimenter’s hand was held over the top of the pole to discourage the mouse from climbing over. Mice were returned to the cage to roam for 30 s between trials. Training was repeated no more than 24 h later (training day 2). For assessment, performed no more than 24 h after the final training day, each mouse was placed head-up at the top of the pole without experimenter intervention and allowed to descend. Each mouse completed three video-recorded trials with 1 min of free roaming between trials. Two measures were analyzed: time-to-descend (TTD), calculated as the mean time from placement until both forepaws contacted the cage floor across the three trials, and time-to-turn (TTT), scored offline from video as the time required for the mouse to rotate into full head-down alignment with the pole. Trials in which a mouse could not turn or descend, fell or slid down the pole, or reversed direction and climbed back up were scored as "physically unable to perform" and recorded in a separate plot; trials in which a mouse descended sideways without fully aligning with the pole were included for TTT but not TTD.

### Cell lines and culture conditions

U937 SLC19A1^KO^ ^20^, U937 SLC19A1^KO^ *LRRC8C*^KO^ ^17^ and HEK293 *LRRC8A*^KO^ ^17^ cells were generated in previous studies. All U937 cell lines were maintained in RPMI supplemented with GlutaMAX, 10% heat-inactivated FBS, and 1% penicillin-streptomycin. HEK 293T and HEK 293 LRRC8A^KO^ were maintained in high glucose DMEM supplemented 10% FBS, 1% penicillin-streptomycin, GlutaMAX, and sodium pyruvate. All cell lines were maintained in a humidified 5% CO_2_ incubator at 37°C, were authenticated by the supplier prior to receipt, and were routinely tested for mycoplasma contamination.

### Reagents

2′3′-cyclic-GMP-AMP (2′3′-cGAMP) was synthesized and purified in-house as previously described^20^ and used for all experiments with cell lines. For primary cell experiments, 2′3′-cyclic-GMP-AMP (2′3′-cGAMP), 2′3′-bisphosphorothioate-cyclic-GMP-AMP (2′3′-cG^S^A^S^MP), 2′3′-cyclic-di-AMP (2′3′-CDA), 2′3′-bisphosphorothioate-cyclic-di-AMP (2′3′-CDAS /ADU-S100), 3′3′-cyclic-GMP-AMP (3′3′-cGAMP), 3′3′-cyclic-di-AMP (3′3′-CDA), and 3′3′-cyclic-di-GMP (3′3′-CDG) were purchased from InvivoGen and reconstituted in endotoxin-free water. STF-1623 was synthesized and purified as previously described^53^.

### Synthesis and purification of ^32^[P]-cGAMP

Radiolabeled ^32^P-ATP (3000 Ci/mmol, Revvityr #BLU002A250UC) was used for synthesis of ^32^P-cGAMP. 1 μM purified sscGAS was incubated with 20 mM Tris-HCl pH 7.4, 2 mM ATP, 2 mM GTP, 20 mM MgCl2, and 100 μg/mL herring testis DNA (Sigma) for 1–3 days. Resulting ^32^P-cGAMP was purified via preparatory TLC as previously described.^20^

### ^32^[P]-cGAMP degradation TLC assay

Tissue dry weight was recorded before samples were transferred into impact-resistant screw-cap tubes (USA Scientific, #1420-9600) and snap frozen in liquid nitrogen. Samples were thawed on ice and topped with 1 ml cold hypotonic lysis buffer (10 mM Tris pH 7.5, 150 mM sucrose including EDTA-free cOmplete protease inhibitor (Roche)) and ∼10 disruption beads. Tissue homogenization was performed using a bead mill homogenizer (Fisher Scientific, #15-340-163) under the following settings: speed 4.5, 30 s, dwell 20 s, 5 cycles. Following homogenization, NaCl was supplemented to 150 mM and NP-40 to 0.1–0.2%. Following a 1h incubation with rotation at 4°C, homogenates were centrifuged at 21,300 × g for 20 min at 4°C, and the supernatants were collected. All concentrations were normalized to 50 mg/ml before testing for the presence of cGAMP hydrolysis. Each CNS region was aliquoted into a PCR tube containing 4 µl lysate, ∼1 nM ^32^P-cGAMP, 5 μM cold cGAMP, physiological levels of ions (15 μM ZnCl_2_, 2.5 mM CaCl_2_, 1 mM MgCl_2_, 5 mM KCl, 100 nM MnCl_2_, 150 mM NaCl), 50 mM Tris pH 7.5, and incubated from 1–24h.

At the indicated timepoints, 1-2 μL from the reaction was spotted on a silica TLC plate (Millipore, Cat# 1.05548.0001) and allowed to dry for a minimum of 15 min. The TLC plates were developed in a mobile phase containing 85% ethanol and 5 mM NH_4_HCO_3._ Plates were dried and exposed to a Storage Phosphor Screen (Cytiva) overnight before being imaged with an Amersham Typhoon 5 Imager (Cytiva). The screens were imaged using a Typhoon 9400 scanner, and the [32]P signal was quantified using Fiji.

### 2’3’-cGAMP quantification

For cGAMP measurements in tissue, hemibrains and spinal cords were excised, placed into impact-resistant screw-cap tubes (USA Scientific, #1420-9600) containing lysis buffer (500 µl for hemibrains or 200 µl for spinal cords) composed of T-PER Tissue Protein Extraction Reagent (Thermo Fisher, #78510) supplemented with ENPP1 inhibitor STF-1623 (1 µM). Tissue homogenization was performed at 4°C using a bead mill homogenizer (Fisher Scientific, #15-340-163) under the following settings: speed 4.5, 30 s, dwell 20 s, 5 cycles. Homogenates were centrifuged at 21,300 × g for 20 min at 4 °C. The supernatants were collected, boiled at 80°C for 10 min to release STING-bound cGAMP,^69^ and then cooled on ice. Supernatant was again collected and used to quantify cGAMP levels using a commercially available ELISA kit (Cayman, #501700) according to the manufacturer’s instructions. When necessary, lysates were diluted to ensure measurements fell within the linear range of the assay. To rule out any interference coming from tissue samples, ELISA standard curves were generated by using brain lysates of healthy, young mice (10 w.k.o) that underwent the same experimental processing. All ELISA measurements were further normalized to protein content in each sample by Pierce660 (ThermoFisher, #22660).

For cGAMP uptake experiments, cells were treated with 10 μM 2’3’-cGAMP for 15 minutes in complete DMEM containing ENPP1 inhibitor STF-1623 (1 μM). For measurements of secreted or intracellular cGAMP in CD11b^+^ microglial and/or myeloid cell culture media, cells were seeded at a density of 2x10^5^ cells/well in 24-well plates after MACS, and cultured in complete DMEM containing STF-1623 (1 μM). After 48h, supernatant was collected, spun down at 21,300xg to remove any cell debris, and used for ELISA. Remaining cells were washed with PBS and lysed in M-PER Extraction Reagent (ThermoFisher, #78501). Extracellular and intracellular cGAMP were quantified using a commercial ELISA kit (Cayman, #501700) following the manufacturer’s instructions and normalized to protein concentrations in the cell lysate.

### 2*’*3*’*-cGAMP stimulation experiments for western blot analysis

For experiments involving inducible U937 cell lines, cells were either left uninduced or induced with 2 µg / mL doxycycline from 24–72 h at a density of 5 x 10^5^ cells / ml before treatment. For experiments involving transient transfection, cells were transfected 24 h before indicated treatment. For all treatments, cells were treated with 50–100 µM of 2*’*3*’**-***cGAMP for 2 h or the indicated concentrations of cyclic dinucleotides. For experiments where cells were treated with glutaminase inhibitor telaglenastat (MedChemExpress, #HY-12248), cells were pre-incubated for 30 min. For experiments where cells were treated with DCPIB (20 µM), L-glutamate (1-2 mM), erastin (50 µM), L-cystine (1 mM), or WAY-213613 (20 µM), cells were pre-incubated for 15 minutes with indicated molecule, then treated with 100 µM 2′3′-cGAMP for 2 h. For experiments involving treatment of U937 cells with glutamine, RPMI without glutamine and containing dialyzed FBS was used. After treatment, cells were pelleted, washed with PBS, and lysed with 50 μL Laemmli Sample Buffer.

### Western blotting

Cell lysates were resolved by SDS–PAGE using precast gels (GenScript, #M42015) and transferred onto nitrocellulose membranes using a dry transfer system (Bio-Rad, #1704150). Membranes were blocked and incubated overnight at 4 °C with primary antibodies diluted 1:1,000-1:4,000, including antibodies against tubulin DM1A (Sigma, CP06), STING (CST, #13647), phospho-STING (CST, #72971), phospho-IRF3 (CST, #4947), Flag (CST, #14793), SLC38A2/SNAT2 (Proteintech 25928-1-AP).

Membranes were washed three times in TBS-T (1× TBS containing 0.1% Tween-20) and incubated for 1 h at room temperature with IRDye-conjugated secondary antibodies (IRDye 800CW goat anti-rabbit, LI-COR, #926-32211; IRDye 680RD goat anti-mouse, LI-COR, #926-68070). After three additional washes in TBS-T, immunoreactive bands were visualized using a LI-COR Odyssey infrared imaging system.

### Electroporation of 2*’*3*’*-cGAMP

U937 SLC19A1^KO^ dox-inducible cell lines were pelleted and resuspended in nucleofector solution (90 mM Na_2_HPO4, 90 mM NaH_2_PO4, 5 mM KCl, 10 mM MgCl_2_, 10 mM sodium succinate) with 100 nM 2’3’- cGAMP to a density of 1 × 10^6^ cells/mL. A volume of 100 μL of cells was then transferred to a 0.2 cm electroporation cuvette and electroporated with program U-013 on a Lonza Nucleofector IIb device. Immediately after nucleofection, 500 μl of warm media was added to the cells. Cells were then transferred to a 24-well plate containing an additional 500 μl of media and incubated at 37°C for 2 h. Following this, cells were collected, lysed with Laemmli Sample Buffer, and resolved by SDS–PAGE for Western blot analysis.

### Generation of Doxycycline Inducible Cell Lines

Lentiviral packaging plasmids (pHDM-G, pHDM-Hgmp2, pHDM-tat1b, and RC/CMV-rev1b) were used to generate lentivirus. Briefly, 500 ng of lentiviral plasmid encoding doxycycline inducible transporters and 500 ng of each of the packaging plasmids were transfected into HEK 293T cells with FuGENE 6 transfection reagent (Promega). Cell supernatant containing lentivirus was harvested after 48 h, and passed through a 0.45 μm filter. To create the inducible cell lines in U937 *SLC19A1*^KO^, 1 mL of filtered supernatant was supplemented with 8 µg/mL Polybrene (Sigma-Aldrich) and added to 5 × 10^5^ cells in a 12-well plate. The next day, the virus containing media was removed and cells were resuspended in fresh media. Cells were selected 48 h later with 10 μg/ml Blasticidin or 400 ng/µl Hygromycin B (ThermoFisher Scientific, #10687010) alongside control cells (uninfected) until all control cells died.

### Primary Microglia and Astrocyte Cultures

Mixed microglia and astrocyte co-cultures were grown as previously reported.^54^ Briefly, the whole brain from adult mice (> 10 w.k.o.) was removed and minced into small pieces. The brain pieces were further dissociated using the Adult Brain Dissociation Kit (Miltenyi Biotec, 130-092-628) according to the manufacturer’s instructions. Dissociated cells were filtered through a 40 µm cell strainer. Myelin was removed by gradient centrifugation using the Debris Removal Kit (Miltenyi Biotec, 130-109-398). Mixed microglia and astrocyte co-cultures were grown in DMEM supplemented with 10% heat-inactivated FBS, 1% penicillin-streptomycin, GlutaMAX, and sodium pyruvate. If cultures were destined for microglia purification, they were additionally supplemented with mouse M-CSF (100 ng/ml, PeproTech, #315-02) and mouse GM-CSF (50 ng/ml, PeproTech, #315-03). 10–12 days after culture was started, cells were magnetically enriched for either microglia or astrocytes.

Before magnetic sorting, conditioned media was collected, cells were lifted with Cell Dissociation Buffer, enzyme free (ThermoFisher, #13151014), washed in 1x MACS buffer (PBS supplemented with 0.5% BSA and 2 mM EDTA), and blocked with FcR Blocking Reagent (Miltenyi Biotec, #130-092-575). For microglia enrichment, cells were labelled with anti-CD11b-beads (Miltenyi Biotec, 130-093-634) and positively selected by magnetic sorting with LS columns (Miltenyi Biotec, 130-042-401) to enrich for CD11b^+^ microglia. For astrocyte enrichment, cells were first labelled with anti-CD11b-beads (Miltenyi Biotec, 130-093-634), and selected by magnetic sorting with LS columns. Negative fraction was collected, labeled with anti-ACSA-2-beads (130-097-678), and positively selected by magnetic sorting. The CD11b^-^ ACSA-2^+^ positive fraction was used for astrocyte culture. Purity of isolated cells was confirmed by flow cytometry to be >90%. Cells were seeded in 1:1 conditioned media:fresh media at a density of 5 × 10^5^ cells/well of 12-well plates and left undisturbed for at least 3 days before being used for any experiment.

### Primary Neuron Cultures

Primary neuron cultures were prepared from cerebral cortices and striata of E18.5–P1 C57BL/6 mice. In brief, cortices and striata were micro-dissected on ice, meninges were removed, and tissue was dissociated with Papain Dissociation System (Worthington Biochemical Corporation, #LK003153) to generate single cell suspensions. Cells were seeded at 300,000–500,000 per well of a 24-well plate pre-coated with Matrigel 24h before seeding. During the first day ex vivo, cells were cultured in MEM supplemented with 10% FBS, GlutaMAX, and glucose. 24h after seeding, the media was replaced with Neurobasal medium (Invitrogen, cat. no. 21103-049) supplemented with B-27 (Invitrogen, cat. no. 17504044), 1% penicillin-streptomycin, GlutaMAX, and. 24 h later, a half media changed was perform to supplement the cultures with 4 µM AraC. Cells were left undisturbed at 37 °C in a humidified chamber of 95% air and 5% CO_2_. Cultures were used for experiments from day 13–40 days after seeding.

### T cell isolation

To isolate homeostatic CD3^+^ T cells as a control for encephalitogenic T cell mtDNA leakage, adult mouse spleens were isolated and mashed through 70 µm cell strainer using the plunger end of a syringe. Cells were washed twice with PBS supplemented with 5 mM EDTA. Red blood cells were lysed in 10 ml ACK lysing buffer (Life Technologies, A1049201) for 10 min at room temperature, inactivated by 40 ml complete medium, and washed twice with PBS supplemented with 5 mM EDTA before staining for FACS.

### Flow cytometry and FACS

Cells were stained for 15 min with 1:1000 LIVE/DEAD™ Fixable Blue Dead Cell Stain (Invitrogen, #L34962) in PBS containing 2 mM EDTA. Following 3 washes with PBS, cells were re-suspended in FACS buffer (PBS supplemented with 2% heat-inactivated FBS and 2 mM EDTA). Before being added to cells, antibodies Fcs were blocked with TruStain FcX (anti-mouse CD16/32) blocking reagent (Biolegend, 101319). Cells were stained with the following antibodies at the indicated dilutions for 20 min at 4°C:

After staining, cells were either analyzed on Attune NxT flow cytometer (ThermoFisher) or sorted using a BDFACS Melody instrument.

### Bulk RNA-seq and data analysis

Cell and tissue RNA was submitted for Plasmidsaurus RNA-Seq (3′ end counting, Illumina NovaSeq X Plus). Gene-level quantification and differential gene expression analysis were performed using the Plasmidsaurus RNA-Seq analysis pipeline, using the mouse Ensembl genome as reference.

### Reverse-transcriptase qPCR (RT-qPCR)

Total RNA was extracted (RNEasyPlus Mini or Micro kit, Qiagen) and quantified with a NanoDrop system. RNA quality was assessed by an absorbance ratio of 260:280 nm. Reverse transcription was carried out with the iScript Reverse Transcription Supermix (Bio-Rad 1708841). qPCR was performed using the LightCycler 480 II (Roche, Basel, Switzerland) with iTaq™ Universal SYBR Green Supermix (Bio-Rad 1725121) and primers listed in table below. Gene expression levels were normalized to GAPDH (unless otherwise specified) and calculated using the 2^-ΔΔCt^ method.

### Subcellular fractionation

Primary cells were grown to 50–80% confluency in 12-well plates or sorted directly into 50 µl of cold 2X digitonin lysis buffer (300 mM NaCl, 100mM HEPES and 50 µg/ml digitonin). At the time of the experiment, cells were washed once with 500 µl ice cold PBS, and lysed with 50 µl cold 1X digitonin lysis buffer (150mM NaCl, 50mM HEPES and 25µg/ml digitonin) for 10 min on ice. Cells were spun at 20,000 xg for 10 min to separate the cytosolic fraction (supernatant) from the heavy membrane organelles and nuclei fractions (pellet). 20 µl of supernatant were carefully removed so as not to disturb the pellet and transferred to a new 1.5 ml microfuge tube, labeled "Cytosolic Fraction". The remaining supernatant was discarded and the pellet was washed once with 500 µl cold PBS. The cytosolic fraction was either diluted 1:10 in MilliQ water for use as a template in qPCR immediately or frozen in -20°C until ready to be assayed. After the pellet was washed, all steps proceeded at room temperature. DNA from the heavy fraction was cleaned up with the DNAEasy Blood and Tissue DNA purification Kit (Qiagen, #69504), using Zymo Spin I Columns (Zymo #C1003-50). Briefly, pellet was resuspended in 200 µl PBS, containing 20 µl Proteinase K (ThermoFisher, #EO0491), and 200 µl AL Lysis Buffer. Samples were incubated at 56°C for 10 min, mixed 200 µl 100% ethanol and loaded into the spin column, washed sequentially with buffers AW1 and AW2, and eluted with 20 µl of Buffer AE. The eluate was labeled "Heavy Fraction" and used at a 1:10 dilution in MilliQ water as a template for qPCR.

### Recombinant DNA and cloning

Plasmid containing the coding sequence (CDS) of human SLC38A2 (Addgene plasmid # 132059 ; http://n2t.net/addgene:132059; RRID:Addgene_132059) and SLC1A2 (Addgene plasmid # 131901 ; http://n2t.net/addgene:131901 ; RRID:Addgene_131901) were obtained from Addgene, deposited by the RESOLUTE Consortium & Giulio Superti-Furga^55^. Custom plasmids (pcDNA3.1(+)) containing the CDS of human SLC7A11 and human SLC3A2 were synthesized by Twist Bioscience. To generate doxycycline-inducible lentiviral plasmids, transporter CDSs were PCR-amplified from the respective parental plasmids and cloned into EcoRI/BamHI-linearized pLVX-TetOne-Flag-Blast or pLVX-TetOne-Flag-Hygro by isothermal Gibson assembly (HiFi DNA Master Mix, NEB E2621). All constructs were verified by full-length plasmid sequencing by Quintara Biosciences.

### Quantification and statistical analysis

All statistical analyses were performed using GraphPad Prism 10 (GraphPad Software). Unless otherwise noted in the figure legends, comparisons between two groups were performed using an unpaired two-tailed Student’s t test with Welch’s correction. Throughout the figures, statistical significance is denoted as follows: ns, not significant (p ≥ 0.05); ∗p < 0.05; ∗∗p < 0.01; ∗∗∗p < 0.001; ∗∗∗∗p < 0.0001. For Western blot quantification, band densitometry was performed using ImageJ/Fiji.

### Single cell RNA-seq and data analysis

#### Tissue processing for single-cell RNA-seq

Animals were deeply anesthetized with isoflurane and transcardially perfused with 10 ml ice cold PBS supplemented with 2 mM EDTA. Following decapitation, four brain regions were isolated and pooled from both hemispheres: cortex, hippocampus, striatum, and diencephalon (thalamus and hypothalamus).

Tissues were finely minced with razor blades and processed with Papain Dissociation System (Worthington Biochemical Corporation, #LK003153) to generate single cell suspensions. Briefly, papain (100 U) was pre-activated by adding 5 ml EBSS and incubated in a 37°C water bath for 10 min. DNAse I (2000 U) dissolved in 500 µl EBSS and 250 µl of was added to each papain tube to generate the digestion mix (20 units/ml papain and 0.005% DNase in EBSS).

Tissues were placed in a 5 ml tube (Eppendorf, #0030119487), containing 800–1000 µl of the digestion mix, and incubated at room temperature without shaking for 30–45 min. Mixture was triturated with a P1000 and any pieces of undissociated tissue remaining after trituration were allowed to settle to the bottom of the tube. Cloudy cell suspension was carefully removed, placed in a clean 1.5 ml microfuge tube, and centrifuged at 300 xg for 5 minutes at room temperature. Supernatant was removed and cell pellet was re-suspended in 800 µl a 1:10 dilution of reconstituted albumin-ovomucoid inhibitor solution in EBSS, containing 0.005% DNase I. Samples were centrifuged at 300 xg for 5 minutes at room temperature and supernatant was discarded.

To remove myelin debris, cell pellets were resuspended to a total volume of 1.55 ml in MACS buffer (PBS supplemented with 0.5% BSA and 2 mM EDTA) and mixed with 450 µl of Debris Removal Solution (Miltenyi Biotec, #130-109-398). 2 ml of MACS buffer were carefully layered on top of this mixture and the solution was spun at 3,000 xg for 10 min, with full acceleration and full brake. After centrifugation, the mixture consisted of three phases (MACS buffer, myelin layer, and cell layer, from up to bottom), of which the top two were aspirated. The remaining phase was mixed with MACS buffer to fill volume by gentle inversion and centrifuged at 4°C and 1000 xg for 10 minutes with full acceleration and full brake.

Supernatant was aspirated and cell pellets were resuspended in 100 µl PBS for cell counting with ViaStain™ AOPI staining solution (Revvity, #CS2-0106-5ML). Single-cell suspensions were loaded onto a Chromium GEM-X 3’ Chip targeting 20,000 cells per sample, and 3′ gene expression libraries were prepared using the Chromium Single Cell 3′ Reagent Kit v4 (10x Genomics) according to the manufacturer’s instructions. Library quality was assessed on an Agilent Bioanalyzer High Sensitivity DNA chip and libraries were sequenced on an llumina NovaSeq X Plus to a depth of 25,000 read pairs per cell.

#### Single-cell RNA-seq alignment and quantification

Sequencing reads were aligned to the mouse reference genome and transcriptome (10x Genomics refdata-gex-mm10-2020-A) and UMI counts were quantified with Cell Ranger count (v8.0.1, 10x Genomics)^56^, including intronic reads (--include-introns=true) with an expected recovery of 20,000 cells per library. All libraries were prepared with Chromium Single Cell 3′ v4 chemistry. Each library corresponded to one brain region (cortex, hippocampus, striatum or diencephalon) from one mouse of each genotype (wild-type, *Enpp1*^H362A^ and *Enpp1*^H362A^ x *Sting*^gt/gt^, giving 12 libraries. Filtered feature–barcode matrices from Cell Ranger were used for all downstream analyses.

#### Quality control and doublet removal

Downstream analyses were performed in Python (3.12.11) with scanpy^57^ (1.11.4) and anndata^58^ (0.12.2). We removed cells with fewer than 300 or more than 6,000 detected genes or with ≥15% of UMIs from mitochondrial genes, and removed genes detected in fewer than 10 cells. Doublets were called separately for each library with Scrublet^59^ (0.2.3; expected_doublet_rate = 0.15, min_counts = 2, min_cells = 3, min_gene_variability_pctl = 85, automatic threshold) and removed. Remaining heterotypic doublets were then removed at the cluster level: we discarded subclusters that co-expressed marker panels of three or more lineages (mean marker detection > 0.30 per lineage), along with cells with more than 30,000 UMIs. Ambient RNA was not removed computationally. Instead, we tracked known ambient transcripts (for example, choroid-plexus Ttr and oligodendrocyte Plp1) in non-expressing cell types when interpreting results.

#### Normalization, dimensionality reduction and clustering

Counts were normalized to 10,000 UMIs per cell and log(x + 1)-transformed. These log-normalized values were used for all visualization, signature scoring and differential expression. We selected 3,000 highly variable genes using scanpy ‘*highly_variable_genes*’ (Seurat flavor, library as batch_key), scaled the data (max_value = 10) and ran principal component analysis (top 50 components). We built a k-nearest-neighbor graph (k = 30) on the first 30 principal components, embedded cells with UMAP^60^ and clustered them with the Leiden algorithm^61^ (leidenalg 0.10.2; resolution 0.5). Each library contained a single genotype–region combination, so no batch integration was applied.

#### Cell-type annotation

We scored each cell for curated marker panels with ‘score_genes’ and assigned each Leiden cluster to the panel with the highest mean score. Assignments were checked against cluster markers (‘rank_genes_groups’, Wilcoxon) and marker dot plots (Figure S3A,B). Marker panels for all 13 cell types are listed in Table S4.

Cell-type composition per region and genotype (Figure S3C,D) is reported descriptively.

#### Microglia subclustering and interferon-responsive microglia

We re-embedded microglia on their own highly variable genes (k = 30, Leiden resolution 1.0). We scored the subclusters for five microglia state signatures: homeostatic, disease-associated microglia (DAM), activated response microglia (ARM), interferon-responsive microglia (IRM) and MHC-II (gene lists in Table S4).

The DAM signature is from Keren-Shaul et al. (2017)^62^; the ARM, IRM and MHC-II signatures are from Sala Frigerio et al. (2019).^63^ Subclusters with choroid-plexus or ependymal contamination were removed, leaving 29,550 microglia. For IRM fractions (Figure S3F,G), 95% confidence intervals were computed with the Wilson method, and genotypes were compared within each region with two-sided Fisher’s exact tests with Benjamini–Hochberg (BH) correction.

#### Differential expression

Differential expression was tested for two contrasts in each cell type: Enpp1 effect: *Enpp1*^H362A^ vs wild-type (WT) STING dependence: *Enpp1*^H362A^ x *Sting*^gt/gt^ vs *Enpp1*^H362A^

We used two-sided Wilcoxon rank-sum tests (‘rank_genes_groups’, method = ‘wilcoxon’) on log-normalized expression across all genes, with BH correction across genes. A contrast was tested only if both groups had at least 30 cells. Genes with FDR < 0.05 and |log2 fold change| > 0.25 were called differentially expressed (DEGs). Analyses were run on all regions pooled and on each region separately. For each cell type, results from the four regions were combined with Fisher’s method^64^ (‘scipy.stats.combine_pvalues’) followed by BH correction. A DEG was called “rescued” when it met the DEG threshold in both contrasts with opposite signs, i.e., the *Enpp1*^H362A^ -induced change was reversed in *Enpp1*^H362A^ x *Sting*^gt/gt^ mice (pathway-agnostic rescue analysis; Figures 3G, 4A,B).

#### Gene set enrichment analysis and Hallmark pathway rescue score

We ran pre-ranked GSEA^65^ with GSEApy^66^ (1.1.10; ‘prerank’), ranking all genes by the Wilcoxon z statistic of each contrast. Mouse gene symbols were converted to their human orthologs by uppercasing. Gene sets came from the MSigDB Hallmark collection^67^ (MSigDB_Hallmark_2020), with gene-set sizes of 15–500 and seed = 42. Pooled-region analyses used 1,000 permutations (results were stable at 10,000 permutations and with two seeds), and per-region analyses used 10,000 permutations. For each cell type and Hallmark pathway, we defined a rescue score as NES_ENPP1 × (−NES_STING), where NES_ENPP1 is the normalized enrichment score for *Enpp1*^H362A^ vs wild-type and NES_STING is the score for *Enpp1*^H362A^ *Sting*^gt/gt^ vs *Enpp1*^H362A^. Positive scores mean that a pathway shifted in *Enpp1*^H362A^ mice shifts back when STING is lost (Figure 3F). Cell type–region combinations whose smallest genotype group had fewer than 100 cells are marked. Rescued DEG lists were also tested for over-representation with Enrichr^68^ (GSEApy ‘enrichr’; MSigDB Hallmark 2020, GO Biological Process 2023, KEGG 2021 Human; BH-adjusted P).

#### Gene signature scoring

Signature scores were calculated with scanpy ‘score_genes’ (ctrl_size = 50, n_bins = 25) on log-normalized expression. Each score is the mean expression of the signature genes minus that of an expression-matched random set of control genes. Genes absent from the data were omitted; notably, Mx1 and Mx2 are not annotated in refdata-gex-mm10-2020-A, so the IRM signature was scored on its remaining eight genes. Signatures: MSigDB Hallmark Interferon Gamma Response, converted to mouse gene symbols (Figures 3L,M, S3G) the microglia state signatures described above full gene lists for all gene sets, with the genes present in the data, are provided in Table S4

#### Statistical analysis of single-cell data

Signature scores and single-gene expression were compared between genotypes, or between expression bins within a genotype, with two-sided Mann–Whitney U tests (‘scipy.stats.mannwhitneyu’).

Transporter-binned analyses (Figures 3L,M, S3G): microglia of each genotype were binned by log-normalized expression of Lrrc8a (Figure 3L,M) or Lrrc8c (Figure S3G) into Zero, Low (0–0.5], Medium (0.5–1] and High (> 1). These were merged into Zero + Low (≤ 0.5) and Medium + High (> 0.5) for testing. Three-group comparisons: tested with Kruskal–Wallis tests.

Associations between gene expression and signature scores: assessed with Spearman correlation. Excitatory vs inhibitory neurons: Mann–Whitney U tests, reporting the rank-biserial correlation as effect size.

Per-gene ISG heatmaps (Figures 5K, S5B): two-sided Wilcoxon rank-sum z statistics from ‘rank_genes_groups’ for a 15-gene ISG panel (Isg15, Isg20, Ifit1, Ifit2, Ifit3, Ifitm1, Ifitm3, Oasl1, Rsad2, Cxcl10, Stat1, Irf7, Usp18, Bst2, Oas1a; Figure 5K) and for the top 40 genes by |z| from a curated 124-gene ISG and inflammatory gene list (Figure S5B; Table SX), with BH correction across all genes in each comparison (* FDR < 0.05).

Box plots: show the median and interquartile range, with whiskers at 1.5× IQR.

Each genotype–region combination came from one animal, so all tests treat cells as observations. P values therefore describe cell-level differences, and we interpreted them together with effect sizes and consistency across brain regions.

**Figure S1.**
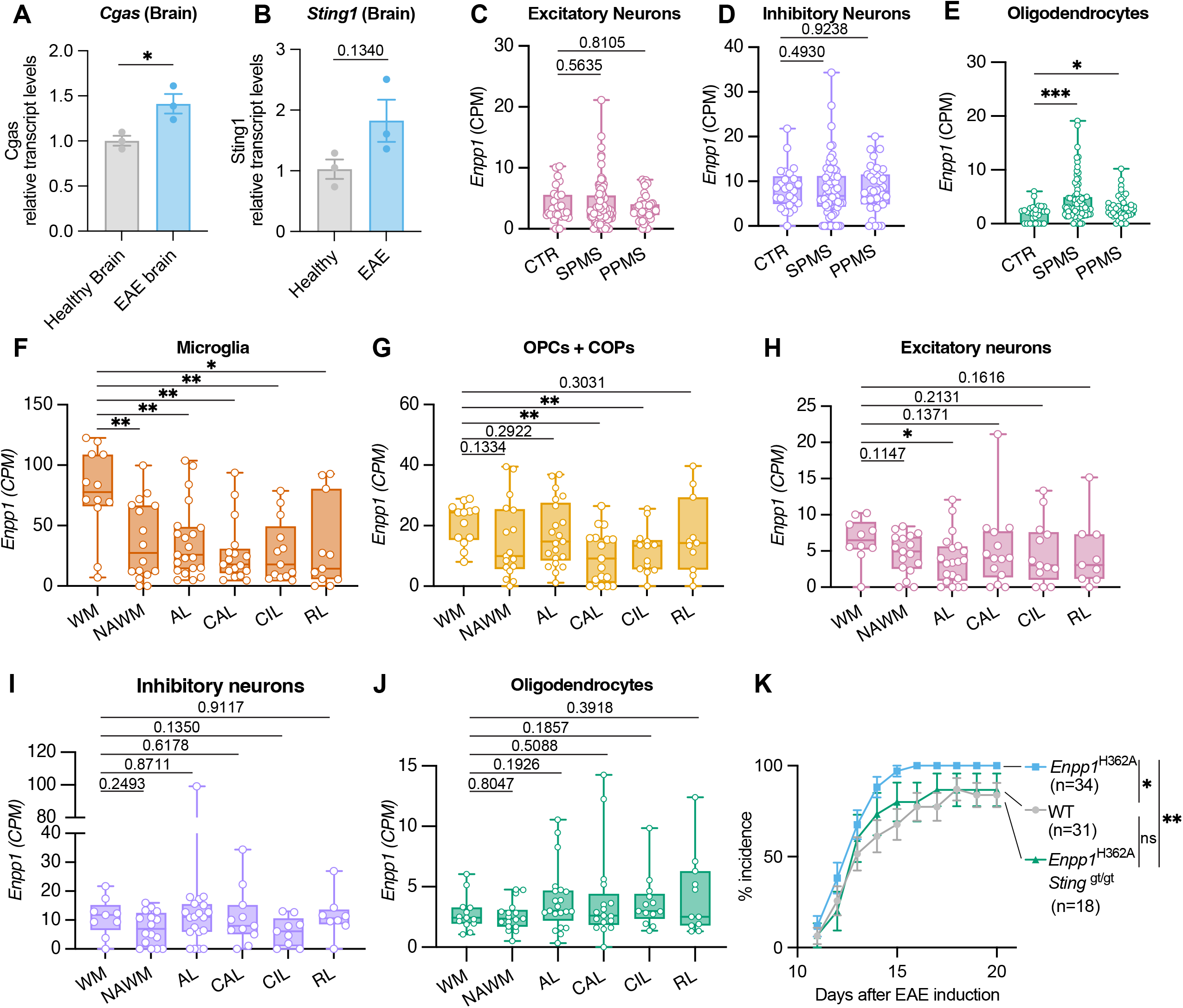
Extracellular cGAMP accumulation from ENPP1 deficiency exacerbates EAE. Related to Figure 1. (A, B) mRNA expression levels of *Cgas* (A) *and Sting* (B) in whole brain tissue of wild-type healthy (n=3 biological replicates) and MOG_35-55_-induced EAE mice (n=3 biological replicates). Data are mean ± s.e.m. *p*-values were calculated using two-sided unpaired t-test with Welch’s correction (C-E) Mean expression of *Enpp1* in excitatory neurons (C), inhibitory neurons (D), and oligodendrocytes (E) in the indicated MS diagnoses. **CTRL**: control; **SPMS**: secondary progressive MS; **PPMS**: primary progressive MS. Significance calculated by Mann-Whitney t-test. (EGA: EGAD00001009169). (F-J) Mean expression of *Enpp1* in microglia (F), oligodendrocyte precursor cells (OPCs) and committed oligodendrocyte precursor cells (COPs) (G), excitatory neurons (H), inhibitory neurons (I), and oligodendrocytes (J) in the indicated white matter MS lesions compared to healthy donors. **WM**: white matter (healthy); **NAWM**: normal-appearing white matter; **AL**: active demyelinated lesion; **CAL**: chronic active demyelinated lesion; **CIL**: chronic inactive demyelinated lesion; **RL**: remyelinated lesion. Data are mean ± s.e.m. *p*-values were calculated using Mann-Whitney t-test. (EGA: EGAD00001009169). (K) Disease incidence in wild-type, *Enpp1*^H362A^ and *Enpp1*^H362A^ *Sting*^gt/gt^ mice at the indicated times after MOG_35-55_ immunization. Significance calculated by two-way ANOVA.

**Figure S2.**
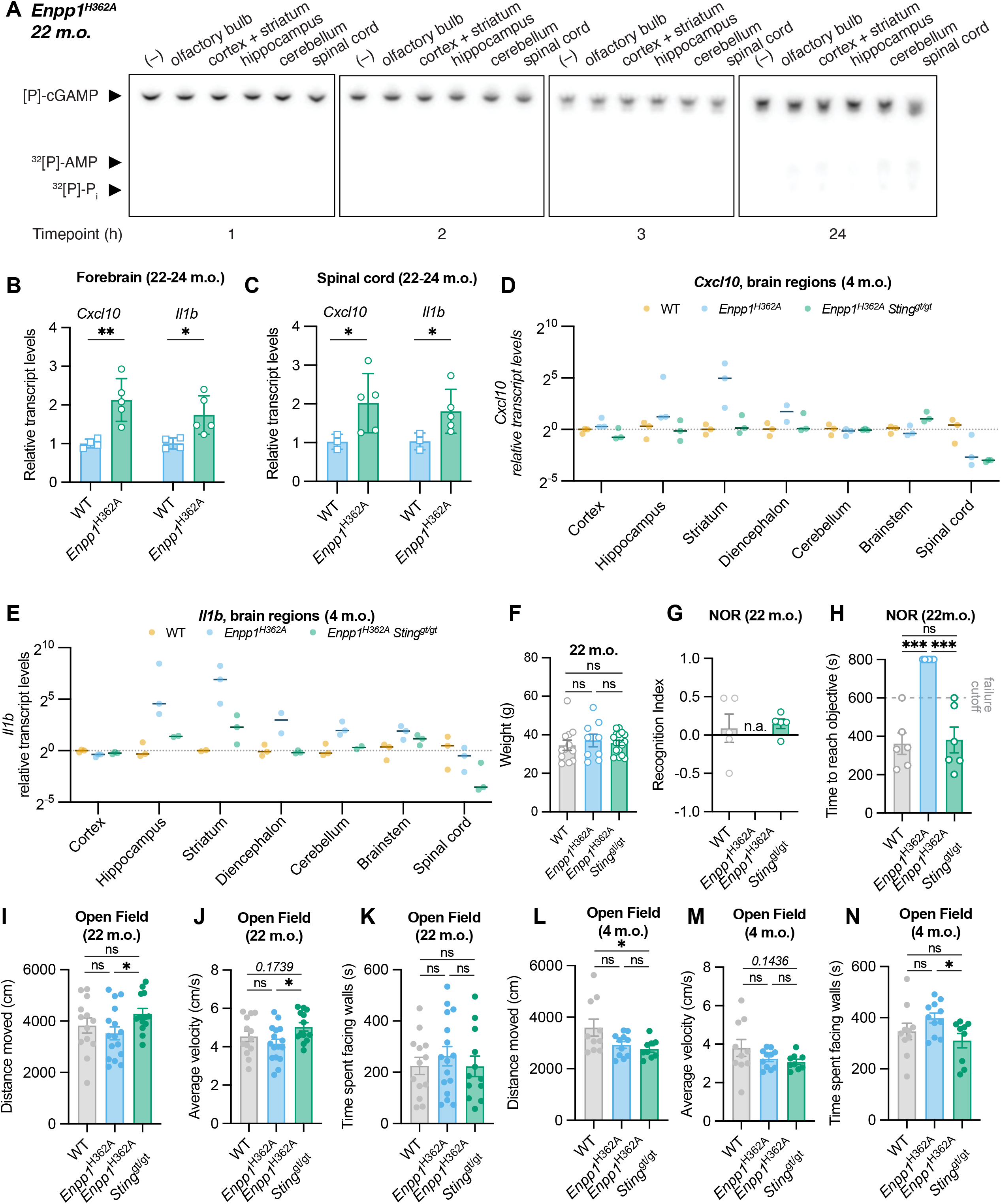
ENPP1-mediated cGAMP hydrolysis is essential for healthy brain aging. Related to Figure 2. (A) Representative thin layer chromatography (TLC) tracking 2′3′-^32^[P]-cGAMP degradation by tissue lysates from the indicated CNS regions of *Enpp1*^H362A^ mice at 22–24 months of age (n=2 biological replicates). (B,C) Relative transcript levels of the indicated ISGs in whole brain tissue (A) and spinal cord tissue (B) of WT (n=3–4) and *Enpp1*^H362A^ (n=4–5) mice at 22–24 months of age. (D,E) mRNA expression levels of *Cxcl10* (D) and *Il1b* (E) at the indicated CNS regions of WT, *Enpp1*^H362A^, and *Enpp1*^H362A^ *Sting*^gt/gt^ mice at 4–6 months of age. (F) Body weights of WT, *Enpp1*^H362A^ and *Enpp1*^H362A^ *Sting*^gt/gt^ mice at 22–24 months of age. Only mice weighing between 25–35 g underwent accelerating rotarod and pole testing. (G) Discrimination index during the novel object recognition test (NOR) for WT, *Enpp1*^H362A^ and *Enpp1*^H362A^ *Sting*^gt/gt^ mice (n=6 mice per group). Only successfully completed tests are plotted. *n.a.* stands for not applicable, which we used for mice from the *Enpp1*^H362A^ genotype that did not interact with either object. (H) Time to reach 30 s of total exploration during the novel object recognition test (NOR) for WT, *Enpp1*^H362A^, and *Enpp1*^H362A^ *Sting*^gt/gt^ mice at 22–24 months of age. The failure threshold of 600s denotes the total duration of the experiment, and 800s denotes the total time of recording (n=6 mice per genotype). (I-K) Distance moved (I), average velocity (J), and time spent at walls (K) during open field testing of WT (n=13), *Enpp1*^H362A^ (n=16) and *Enpp1*^H362A^ *Sting*^gt/gt^ (n=13) mice at 22–24 months of age. (L-N) Distance moved (L), average velocity (M), and time spent at walls (N) during open field testing of WT (n=10), *Enpp1*^H362A^ (n=11) and *Enpp1*^H362A^ *Sting*^g/gtt^ (n=9) mice at 22–24 months of age. Data are mean ± s.e.m. *p*-values were calculated using two-sided unpaired t-test with Welch’s correction unless otherwise indicated.

**Figure S3.**
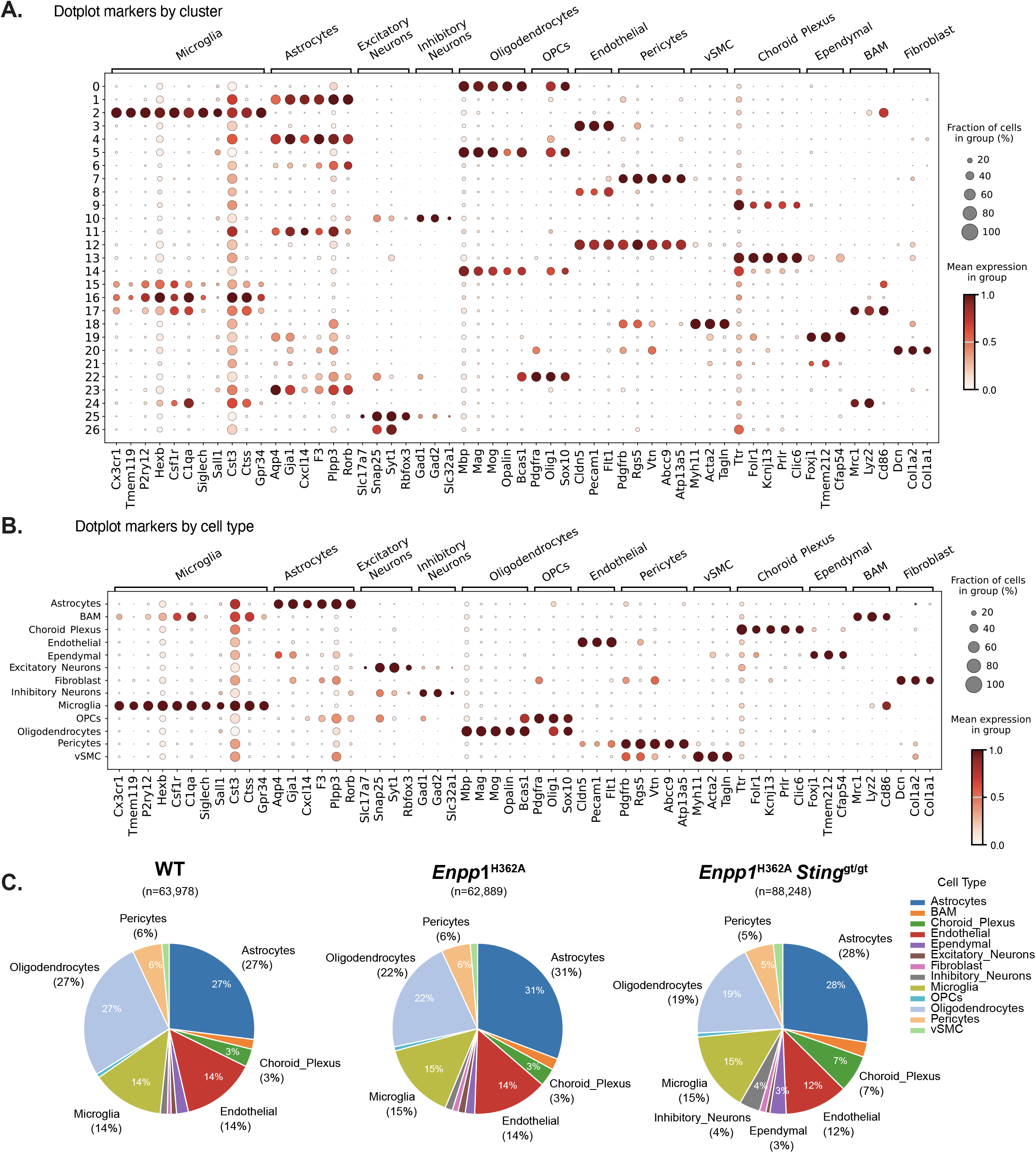
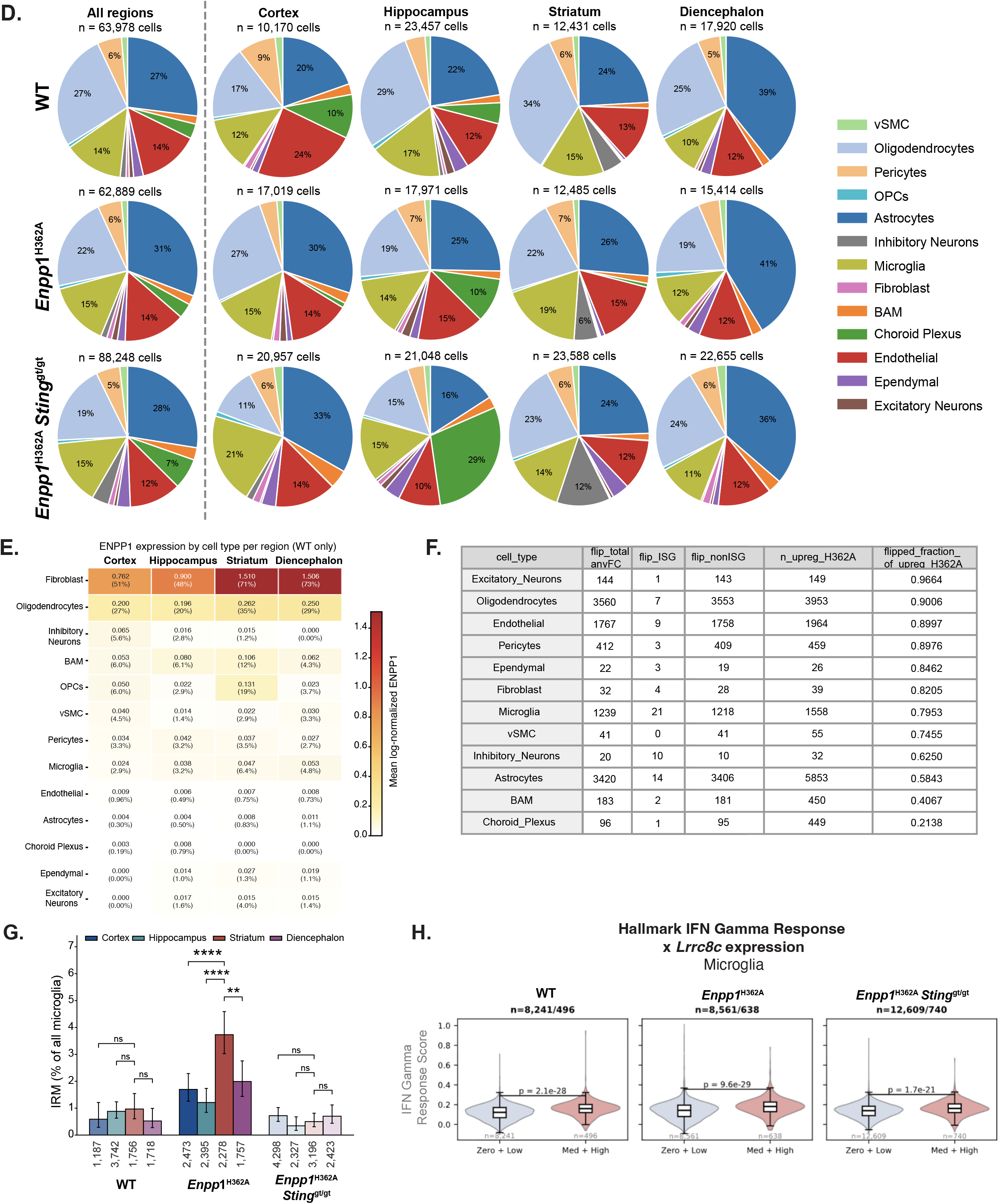
Cellular composition and marker gene expression distribution in scRNAseq. Related to Figure 3. (A,B) Bubble heatmaps showing expression of selected marker genes for cluster annotation (A) and cell type assignment (B). Dot size indicates the fraction of expressing cells, colored based on average expression levels. (C) Pie plot of cell type compositions of pooled brain regions of WT, *Enpp1*^H362A^ and *Enpp1*^H362A^ *Sting*^gt/gt^ mice at 4–6 months of age. (D) Pie plot of cell type compositions across the indicated brain regions of WT, *Enpp1*^H362A^ and *Enpp1*^H362A^ *Sting*^gt/gt^ mice at 4–6 months of age. (E) *Enpp1* expression levels and percent expressing-cell types in different brain regions of wild-type mice at 4–6 months of age. (F) Summary of rescued DEGs in different cell types of 4–6 months old mice (pct_threshold of expressing cells = 0.1). (G) Fraction of interferon-responsive microglia (IRM) by region in wild-type, *Enpp1*^H362A^ and *Enpp1*^H362A^ *Sting*^gt/gt^ mice at 4–6 months of age. (H) Expression of Hallmark IFN-γ response signature in microglia binned by expression of *Lrrc8c*. Hallmark IFN-γ response module score in microglia from wild-type, *Enpp1*^H362A^,and *Enpp1*^H362A^ *Sting*^gt/gt^ mice at 4–6 months of age, all four brain regions pooled. Microglia were binned by log-normalized *Lrrc8c* expression. Inner box plots show the median and interquartile range, with whiskers at 1.5× IQR. P-values were calculated from two-sided Mann–Whitney U tests.

**Figure S4.**
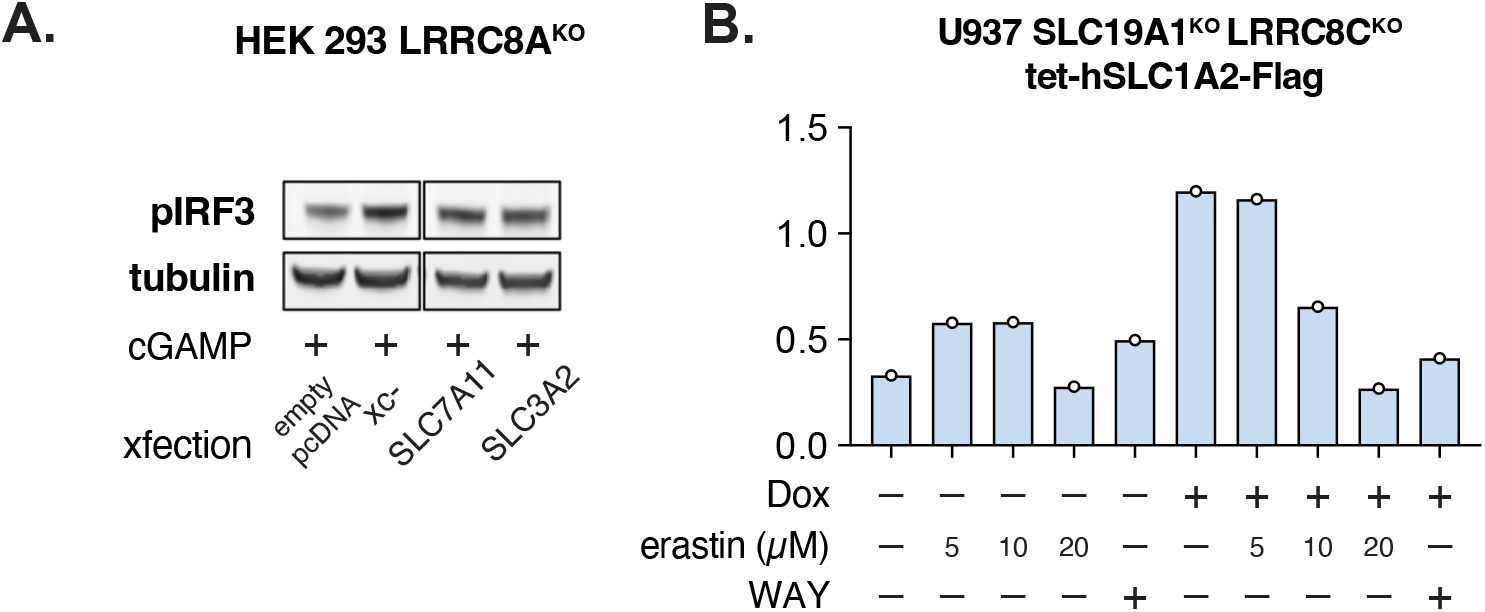
Astrocytes can import extracellular cGAMP through system xc-facilitated by glutamate transporters. Related to Figure 4. (A) STING signaling output as measured by pIRF3 levels in HEK 293 LRRC8A^KO^ cells expressing system xc- (SLC7A11 and SLC3A2), xCT (SLC7A11 alone), or 4F2hc (SLC3A2 alone) after treatment with or without 100 µM 2′3′-cGAMP. One representative Western blot is shown out of n = 2 biological replicates (B) STING signaling output as measured by pIRF3 levels in U937 *SLC19A1*^KO^ *LRRC8A*^KO^ tet-h*SLC1A2*-Flag cells treated with or without 2 µg/ml doxycycline (Dox) for 72 h, then treated with or without 50 µM 2’3’-cGAMPand/or erastin or WAY-213613 (20 µM).

**Figure S5.**
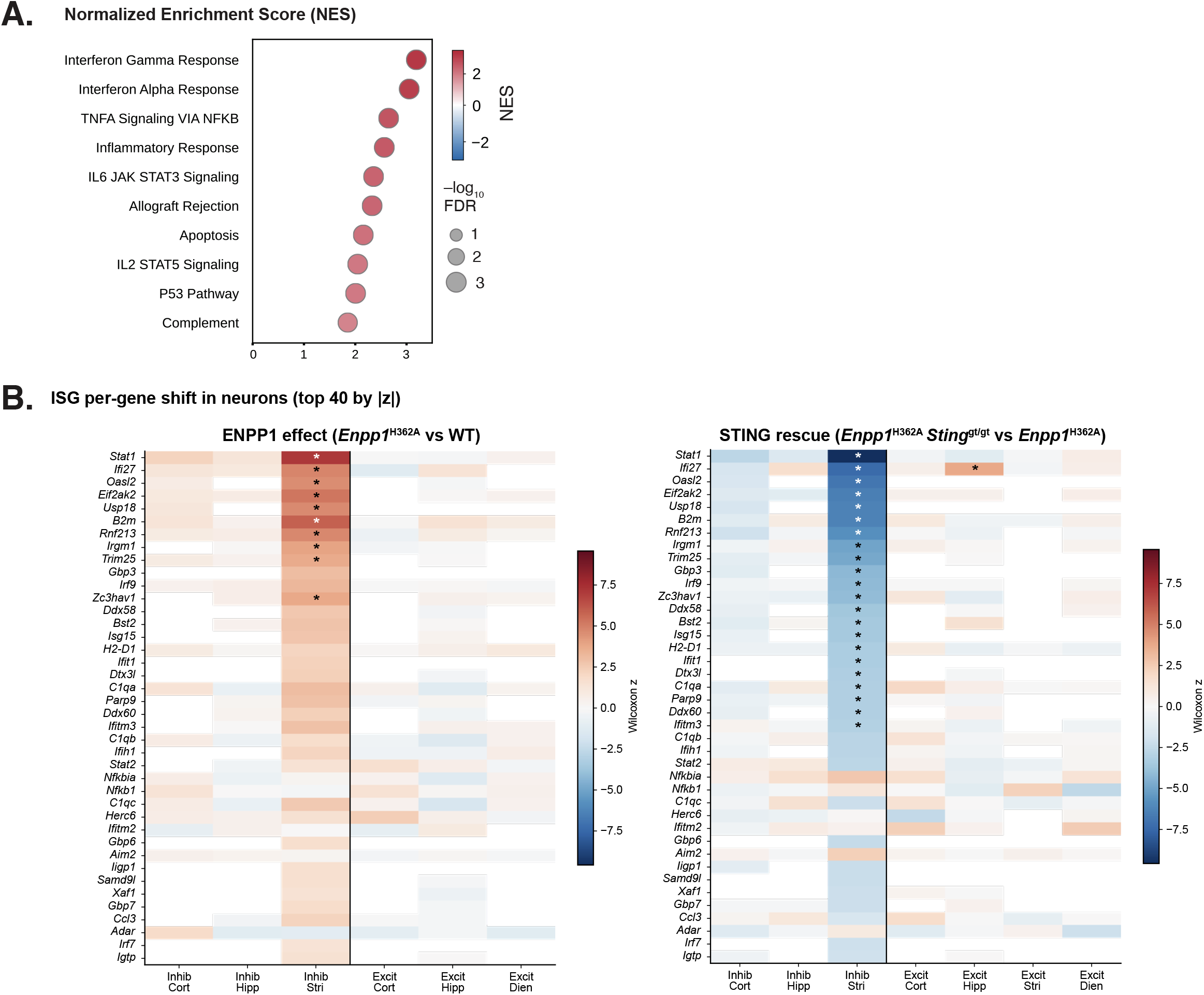
Extracellular cGAMP can activate neuronal STING through system xc^_^ facilitated by glutamine transporters and glutaminase. Related to Figure 5. (A) Functional enrichment sorted on adjusted *p*-value in ascending order. Pathways identified with gene set enrichment analysis (GSEA) using the Hallmark gene set reveal interferon response as the top-enriched pathway in bulk RNA-seq analysis of primary mouse cortical and striatal neurons treated with 10 µM 2′3′-cGAMP and 1 µM ENPP1 inhibitor STF-1623 after 24h. (B) Per-gene ISG responses in inhibitory and excitatory neurons by brain region. Heatmap of the Wilcoxon rank-sum z-score for the 40 ISGs with the largest absolute z-score in any column, selected from a curated 124-gene ISG list (118 detected). Left, *Enpp1*^H362A^ vs. WT; right, *Enpp1*^H362A^ *Sting*^gt/gt^ vs. *Enpp1*^H362A^. Positive (red) z indicates higher expression in the first-named genotype. * FDR < 0.05, Benjamini–Hochberg across all genes in each comparison. Region × neuron-type combinations with fewer than 30 cells per genotype were not tested and are not shown (inhibitory diencephalon; excitatory striatum, left).

**Table S1.** Antibodies used for flow cytometry and FACS.

| Antibody | Clone | Dilution | Supplier | Cat# |
| --- | --- | --- | --- | --- |
| CD45-APC | 53-7.3 | 1:200 | Biolegend | 100625 |
| CD11b-FITC | M1/70 | 1:200 | Biolegend | 101206 |
| CD3-PE | 17A2 | 1:200 | Biolegend | 100206 |
| ACSA-2-PE | IH3-18A3 | 1:100 | Miltenyi Biotec | 130-123-284 |

**Table S2.**
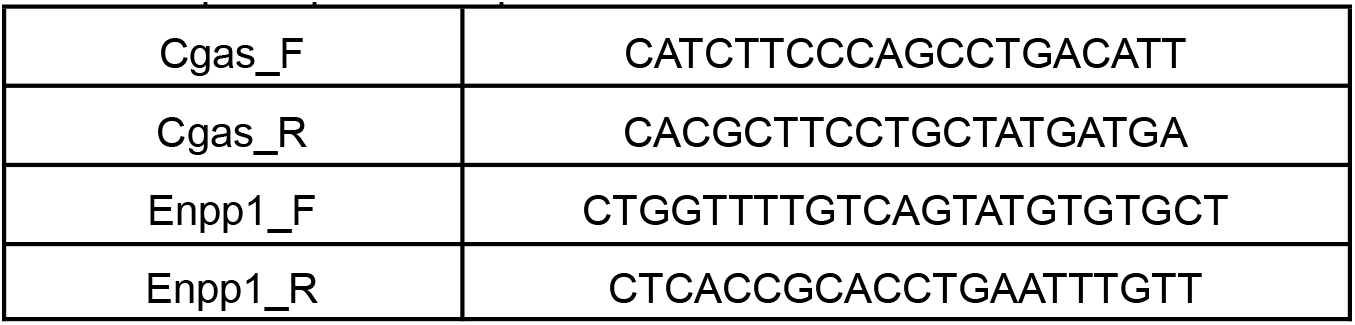
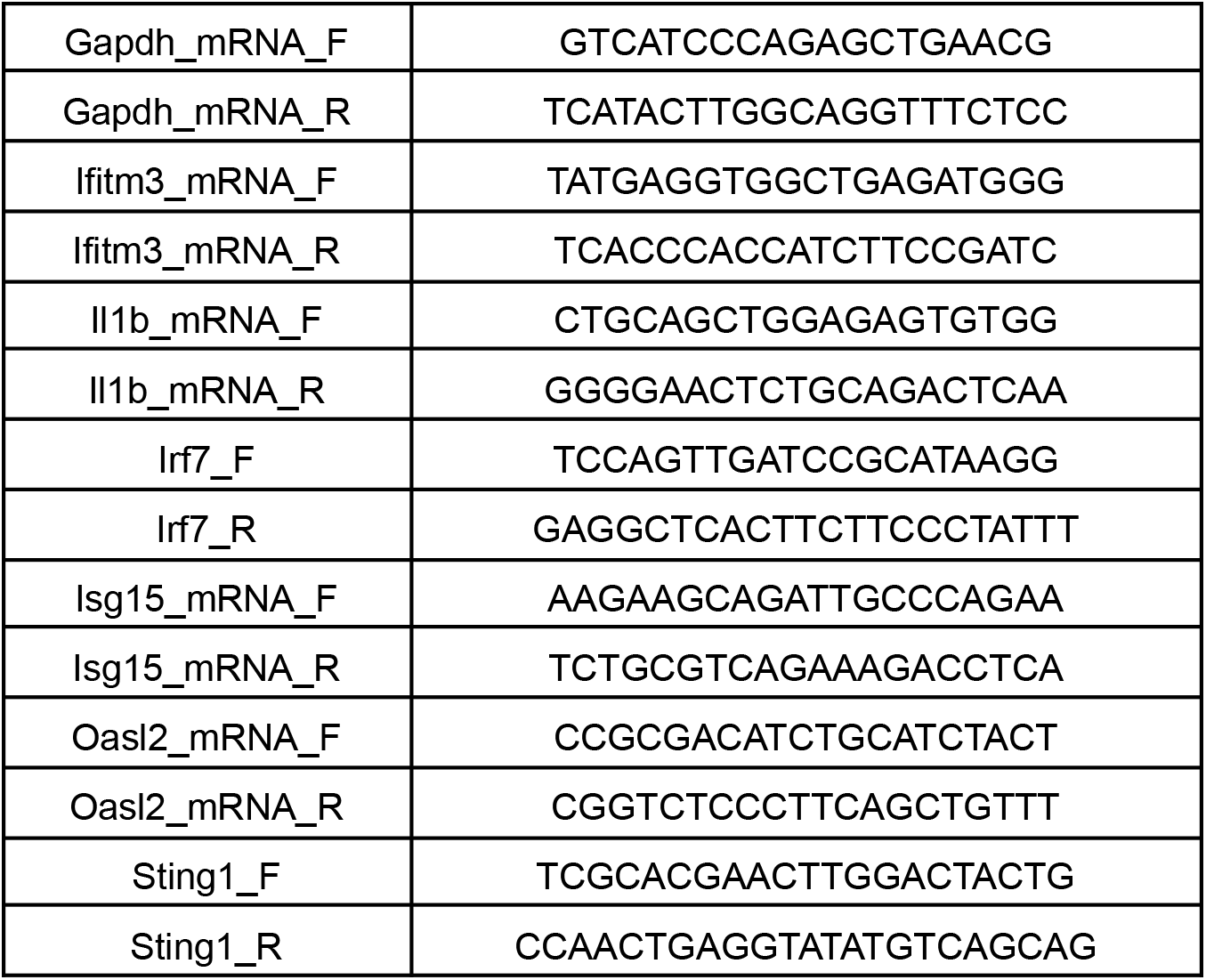
qPCR primer sequences.

**Table S3.**
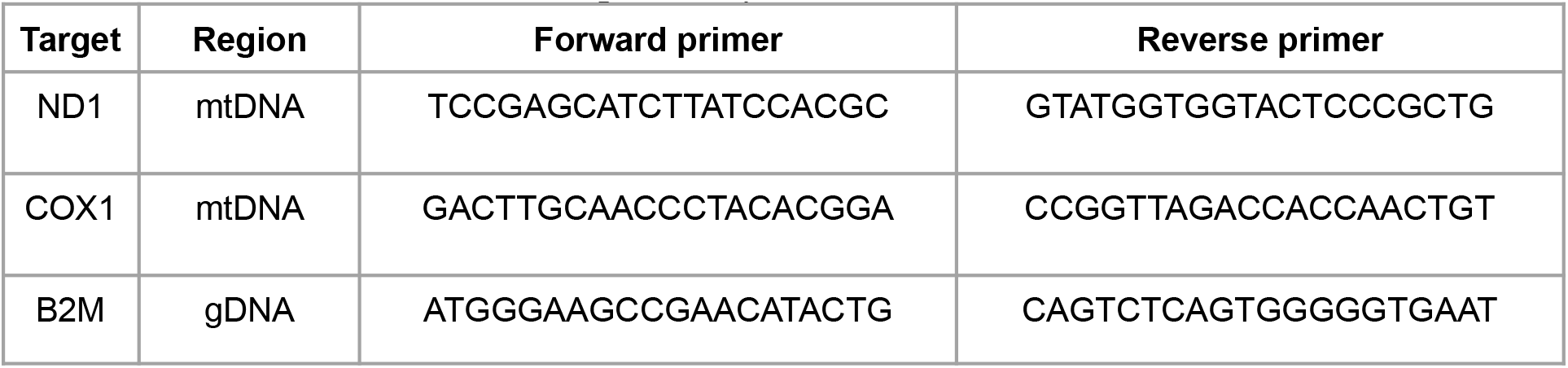
Primers used for mtDNA and gDNA amplification.

